# Targeting of PTP4A3 overexpression sensitises HGSOC cells towards chemotherapeutic drugs

**DOI:** 10.1101/2024.10.28.620719

**Authors:** Ana López-Garza, David James, Emma Creagh, James T. Murray

## Abstract

Of all the gynecologic malignancies, Ovarian cancer (OC) has the highest mortality rate, partly attributable to its propensity for chemotherapy resistance. The most common sub-type of OC is serous, of which High-Grade Serous Ovarian Cancer (HGSOC) is the most lethal sub-type. Elevated expression of Protein Tyrosine Phosphatase 4 A3 (PTP4A3) is implicated in tumour cell invasion and metastasis, by upregulating the PI3K/Akt/mTORC1 axis. Previously we reported PTP4A3 increased the survival of non-serous OC cells *in vitro* by activating the autophagy pathway.

The present study focused on understanding the impact of PTP4A3 on cell growth, proliferation, and autophagy in HGSOC cells. In particular, we sought to understand whether targeting PTP4A3 in cancer cells that overexpress this phosphatase would sensitise HGSOC cells to existing chemotherapeutic drugs. We report that shRNA-mediated gene silencing of PTP4A3 resulted in the upregulation of compensatory mechanisms that may render PTP4A3 targeting redundant as a monotherapy. However, pan-PTP4A inhibition with JMS-053 overcame this. Finally, silencing of PTP4A3 expression sensitized HGSOC cells to clinically relevant chemotherapeutic drugs.

Since mAb therapies targeting PTP4A3 are already in clinical trials, therapeutic targeting of PTP4A3 may have significant value in improving outcomes for those patients with HGSOC, in the clinical setting.

## INTRODUCTION

Ovarian cancer (OC) is the most lethal gynaecologic malignancy and the fifth leading cause of cancer-related deaths among women. Although it is less common than breast cancer, the fatality rate of OC is three times higher, with a 5-year survival rate of 46.5% (1–3). This high mortality rate is attributed to poor early detection diagnostic methods, inadequate therapeutic options, high recurrence rates, and its propensity for drug resistance (1, 2). Due to its limited detection at early stages, four in every five women are diagnosed when the cancer has already metastasized to other tissues (1–3). Serous is the most common sub-type of OC and is subdivided into High- and Low-Grade Serous Ovarian Cancer (HGSOC and LGSOC). Two-thirds of ovarian cancer mortality is specifically due to the HGSOC subtype (2).

The prenylated protein tyrosine phosphatase (PTP), PTP4A3, is one of three PTPs (PTP4A1-3, also known as Phosphatase of Regenerating Liver 1-3) that share >75% sequence identity (4, 5). Ordinarily, PTP4A3 is involved in the early development of the circulatory system, with expression detected in the foetal heart, developing blood vessels and pre-erythrocytes, but not in adult tissues (6). More recently, PTP4A3 expression has been implicated in tumour cell invasion and metastasis through disruption of cell adhesion, while PTP4A3 is transcriptionally up-regulated by p53 (6, 7). PTP4A3 is also involved in the regulation of the epithelial-mesenchymal transition (EMT), through its inhibition of PTEN and activation of PI3K/AKT signalling (6). The enzymatic activity of PTP4A3 is essential for its function since the inactive mutant C104S is unable to activate these pathways (8).

Autophagy is a cellular catabolic process by which cytoplasmic material is ultimately degraded by the lysosome. It functions as a homeostatic cellular recycling mechanism to ensure cellular survival by minimizing the accumulation of cellular damage. Autophagy is initiated when cells are confronted with potentially dangerous environmental, physical, chemical, or metabolic signals, such as thermal stress, irradiation, changes in pH/osmolality, or a shortage in nutrients and/or oxygen (9–11). Dysfunctional autophagy is associated with numerous human diseases, including diabetes, response to infection, neurodegeneration, and cancer (12, 13).

Autophagy plays a complex role in cancer development and progression that depends on the phase of carcinogenesis and the tumour context. Drugs that induce or inhibit autophagy have both been described to have anti-cancer effects (9). Autophagy can limit genomic instability and tumour development by protecting cells from ROS-induced DNA and proteotoxicity (15) and chronic upregulation of autophagy is associated with the induction of cell death (9). In contrast, upregulation of autophagy in growing tumours can compensate for limited nutrient supply and mitigate genotoxic and metabolic stresses. This is underscored in some cancers where the KRAS oncogene can drive autophagy addiction (16–19). The pro-tumourigenic activity of autophagy is also associated with drug resistance and represents a target for combinatorial therapeutic approaches (12, 20). Anti-cancer therapies that target autophagy are dependent on actual levels of ongoing autophagy in tumour cells (21, 22).

PTP4A3 expression is upregulated in multiple human cancers, including OC (9, 23). Over-expression of PTP4A3 in OC has been detected in advanced stage-III when compared to early stage-I tumours, suggesting a correlation between PTP4A3 expression and the invasiveness of OC (24, 25). Our previous study demonstrated that PTP4A3 increases the survival of ovarian cancer cells *in vitro* by activating the autophagy pathway (11). High *PTP4A3* expression was shown to positively correlate with the expression of two critical autophagy genes, *PIK3C3* and *BCLN1,* which significantly predict poorer OC prognosis in patients, suggesting a critical role of autophagy in PTP4A3-driven OC progression (9).

The present study focused on understanding the impact of PTP4A3 on cell growth, proliferation, and autophagy mechanisms in OC cells. In particular, we sought to understand how PTP4A3 regulates autophagy in HGSOC cells, and whether targeting PTP4A3 in cancer cells that overexpress this protein sensitises them to chemotherapeutic drugs.

## MATERIALS AND METHODS

### Cell culture

OVCAR 3, OVCAR 4 and Kuramochi cell lines were kindly provided by Dr Nuala McCabe (The Queen’s University, Belfast). They were maintained in RPMI-1640 (Sigma-Aldrich, UK R8758) supplemented with 10 % (v/v) Foetal Bovine Serum (Sigma-Aldrich F7524), 1 % (w/v) L-glutamine (Sigma-Aldrich G7513) and 1 % (w/v) Penicillin/Streptomycin (Sigma-Aldrich P4333) at 37 °C, 95 % humidity and 5 % CO_2_.

### Stable knockdown cell line generation

Replication-defective lentiviral particles were generated using HEK293T cells PEI transfected with 4.5 µg pCMV-DVPR (packaging plasmid), 1.5 µg pCMV-VSV-G (envelope plasmid) and 6 µg pLKO.1 (TRCN000001883 PTP4A3 MISSION shRNA, or scramble control plasmid, Sigma-Aldrich). Media containing lentiviral particles was harvested after 48 h, filtered through a 0.45 µm sterile filter and then added to target cells. Selection of PTP4A3 silenced Kuramochi and OVCAR 4 cells was performed with 1.6 µg mL^-1^ and 0.8 µg mL^-1^ puromycin, respectively. Knockdown efficiency was validated by Western immunoblot.

### Transient protein overexpression

pEGFPC1 parent plasmids (Clontech) were further cloned in-house to generate PTP4A3 expressing (9). Plasmids were amplified using chemically competent NEB5α *E. coli* cells (New England Biolabs). The cDNA Plasmids were isolated using a HiSpeed Plasmid Maxi Kit (QIAGEN, Manchester, UK). Once eluted, the concentration (A280 nm) and purity (A260/A280 nm) of plasmid DNA were assessed using a NanoDrop spectrophotometer ND-1000. To overexpress a protein of interest, target cells were PEI transfected with 4.6 μg cDNA. Transfection efficiency was assessed by fluorescence microscopy and Western immunoblot.

### Cell growth and death monitoring

Cells were plated in 96-well plates before adding increasing doses of 5- fluorouracil (0-1 mM), cisplatin (0-40 µM), paclitaxel (0-100 nM) or JMS-053 (0- 25 µM) together with 2.5 µg mL^-1^ propidium iodide (PI) to stain dead cells. Plates were transferred to the Incucyte S3 and the system was set to take 4 images per well every 3 h for 3 d. Confluency and the number of PI-positive cells were quantified using the Incucyte software and presented using GraphPad Prism 9 software.

### Western blot analysis

Whole-cell protein lysates were sonicated using a MicroTip sonicator (20 % output, for 10 s) and protein concentration was quantified using a Bradford protein quantification assay (Thermo-Pierce). Lysates were separated by SDS-PAGE, transferred onto PVDF membranes, and incubated in Blocking buffer of 5 % (w/v) non-fat dried milk (NFDM) in Tris-Buffered Saline, 0.1 % (v/v) Tween 20 (TBS-T) for 30 min at room temperature. Membranes were incubated overnight at 4 °C on a rocker with the following antibodies in Blocking buffer: β-actin (Sigma-Aldrich, Gillingham, UK A5441), LC3B (Cell Signaling Technology, Leiden, Netherlands 2775), S6K1 (9202), p-S6K1 (9205), ERK1/2 (9102), p-ERK1/2 (9106), ULK1 (4773), p-ULK1 (6888), p-AKT (9271), PTP4A3 (Santa Cruz Biotechnology, Heidelberg, Germany sc-130355), AKT (26), pan-PTP4A3 (Bio-Techne MAB32191), and GFP (Thermo Fisher Scientific, Horsham, UK A-11122), then washed thrice with TBS-T and incubated with species-appropriate HRP-conjugated secondary antibodies (Thermo Fisher Scientific) diluted in Blocking buffer. After three additional washes in TBS-T, membranes were developed with ECL (Merck Millipore, Gillingham, UK 638173). The intensity of protein bands was quantified by densitometry using ImageLab software and presented using GraphPad Prism 10 software.

### Statistical analysis

GraphPad Prism 10 software was employed for statistical analysis. All data are represented as mean ± S.E.M., with experiments done as n=3, unless otherwise stated in the figure legend. Statistical significance was indicated when p<0.05. A one-way ANOVA test was used to compare two conditions. A two-way ANOVA with a Tukey multiple comparisons test was used to compare more than two groups of individual treatments.

## RESULTS

### PTP4A3 expression and PI3K and Ras activity in HGSOC cells

The expression of PTP4A3 was investigated in three HGSOC cell lines, Kuramochi, OVCAR 4 and OVCAR 3 (27). There was negligible detection of PTP4A3 protein in OVCAR 3 cells, whereas Kuramochi cells express high levels, and OVCAR 4 cells expressed ∼10-fold less PTP4A3 than Kuramochi cells (Fig. 1a, b). We categorised these cell lines as having no (OVCAR 3), low (OVCAR 4) and high (Kuramochi) PTP4A3 expression for subsequent investigations.

**Figure 1.**
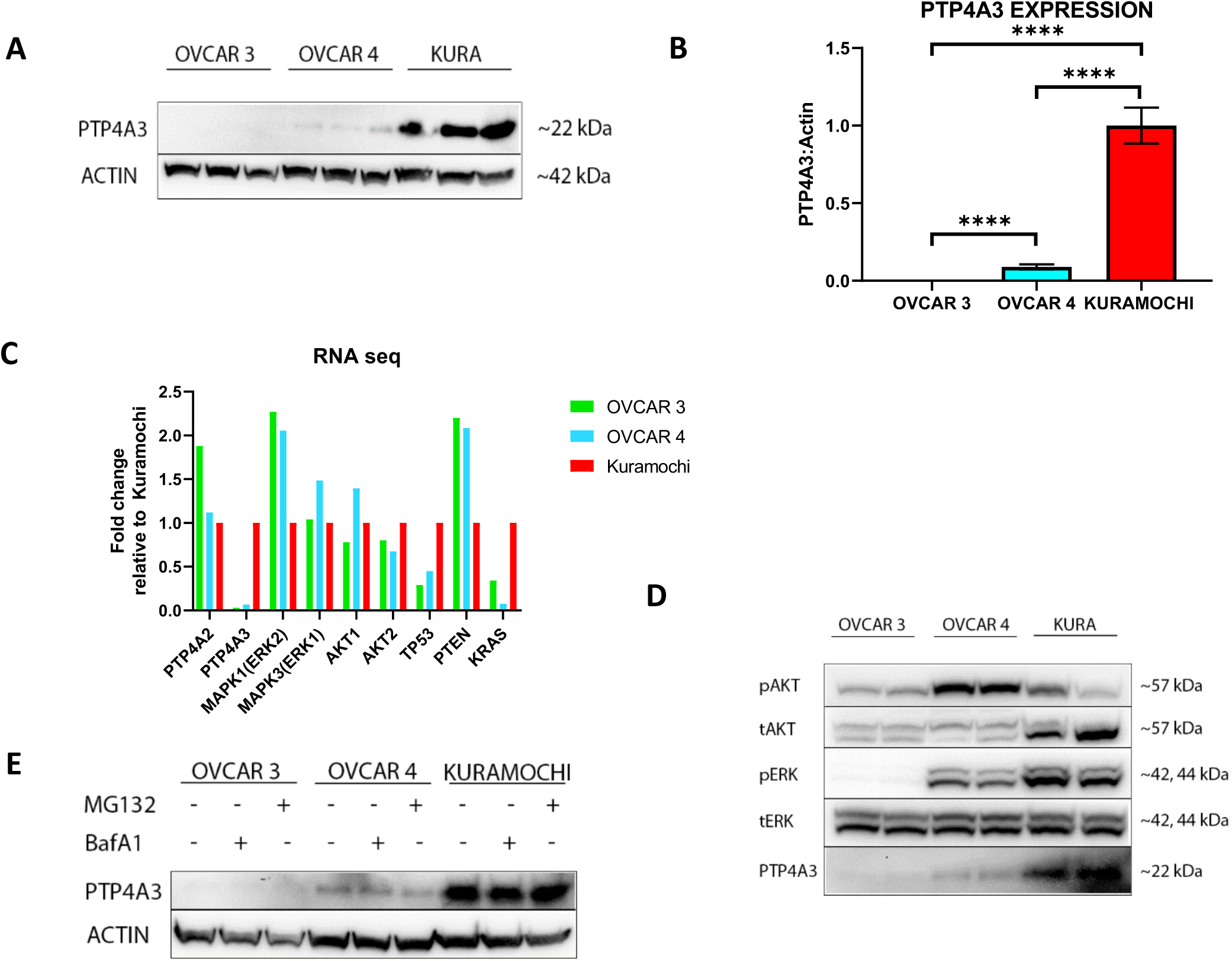
The expression of PTP4A3, PI3K and Ras signalling pathway proteins varies across HGSOC cell lines. (**A**) Expression of PTP4A3 was assessed in OVCAR 3, OVCAR 4 and Kuramochi cells by Western immunoblot and (**B**) this was quantified by densitometry, with PTP4A3 normalised to β-actin. (**C**) NCBI GEO repository RNA-seq data was analysed for gene expression in the three HGSOC cell lines, normalised to Kuramochi levels. (**D**) Cell lysates were analysed for phosphorylation of AKT/PKB (pSer473) and ERK1-2 (pThr202- Tyr204). Antibodies that recognise total AKT/PKB and ERK1-2 were used as controls. Expression of PTP4A3 is shown for comparison. (**E**) Turnover of PTP4A3 was analysed by incubating with or without either MG132 (2.5 µM, 2 h) or BafA1 (100 nM, 1 h) before lysates were collected with β-actin as loading control. Results are representative of at least three independent experiments. Error bars = ± S.E.M. **** p< 0.0001 by unpaired *t*-test.

PTP4A3 is implicated in both RAS and PI3K pathway signalling. Data collected from Cellosaurus and Harmonizome combining CCLE, BioGPS, and GDSC databases revealed that OVCAR 3 cells express mutated PIK3R1 that affects the PI3K signalling pathway, while OVCAR 4 cells harbour mutations in MAP3K1 in the RAS pathway. Moreover, OVCAR 3 and Kuramochi also possess high KRAS mRNA expression, whereas OVCAR 4 cells have low mRNA expression of MEK2, suggesting dysregulation of RAS pathway signalling by opposing mechanisms. Finally, Kuramochi and OVCAR 3 cells have been shown to possess high and low PTP4A3 mRNA expression levels, respectively (28). Analysis of RNA-seq datasets, normalised to Kuramochi expression levels, were compared (Fig. 1c). *PTP4A3* RNA expression in OVCAR 3 and OVCAR 4 was only 3% and 6.7%, respectively, of the amount expressed in Kuramochi, which correlated with PTP4A3 protein levels (Fig. 1a, b). Interestingly, *PTP4A2* gene expression was almost 2-fold higher in OVCAR 3 cells than OVCAR 4 and Kuramochi cells (Fig. 1c). *KRAS* amplification in Kuramochi also observed in the RNA-seq datasets, with OVCAR 3 having 3-fold less and OVCAR 4 having >12- fold less *KRAS* expression than Kuramochi cells. In agreement with previous studies, the expression of *TP53* correlated with PTP4A3 expression levels (29), whereas *PTEN* was inversely correlated with *PTP4A3* expression (24) across the three cell lines (Fig. 1c).

The PI3K/AKT and RAS signalling pathways are strongly involved in HGSOC, and PTP4A3 regulates AKT signalling (6–8). When the basal activity of these pathways was assessed in Figure 1d, we found that KRAS amplification in Kuramochi cells (28) correlated with their elevated levels of ERK phosphorylation. OVCAR 4 cells exhibited high PI3K activity, as indicated by phospho-AKT expression, but lower RAS activity than Kuramochi, in agreement with their KRAS expression status. However, the low KRAS activity in OVCAR 3 cells did not match their elevated KRAS expression, although the lack of PTP4A3 expression may partly explain the low activity of both pathways in these cells (Fig. 1d).

Expression of PTP4A3 may be controlled through proteostatic mechanisms, with exogenously expressed PTP4A3 behaving as an autophagy substrate (9). However, when HGSOC cells were treated with or without the inhibitors bafilomycin A1 (BafA1) or MG132, to inhibit autophagy and the ubiquitin-proteasome systems, respectively, immunoblotting detected no difference in PTP4A3 expression levels in any cell line (Fig.1e).

### Amino acid deprivation-induced autophagy is dysregulated in OVCAR 4 and Kuramochi cells

Given both the paradoxical role of autophagy in cancer and PTP4A3 overexpression promoting autophagy to enhance tumour cell growth, the autophagy status of HGSOC cells was examined (9, 13). To assess basal and inducible autophagy, each cell line was deprived of amino acids (EBSS) for up to 8 h, and the activation of autophagy was quantified by monitoring the conversion of the autophagy marker protein LC3B, from LC3B-I to LC3B-II. The removal of amino acids results in mTORC1 inhibition concomitant with autophagy activation, which was assessed by monitoring S6K1 Thr389 phosphorylation (30–32).

In all three cell lines, there was robust inhibition of mTORC1 activity after 2 h EBSS treatment (Fig. 2a, c, e). OVCAR 4 cells also significantly induced the formation of LC3B-II within 2 h EBSS whereas no statistically significant LC3B-II induction was observed in OVCAR 3 or Kuramochi cells (compare Fig. 2a, d with Fig. 2b, e and Fig. 2c, f). Despite this, there was a rapid loss of LC3B-II in OVCAR 3 cells over an extended time frame (Fig. 2a, d), but this was not evident in Kuramochi cells (Fig. 2c, f). Although OVCAR 4 cells rapidly induced LC3B-II at 2 h following amino acid withdrawal, LC3B-II turnover was minimal and not statistically significant (Fig 2b, e). Finally, Kuramochi cells showed minimal activation of autophagy, with no change in LC3B-II levels following amino acid withdrawal (Fig. 2c, f).

**Figure 2.**
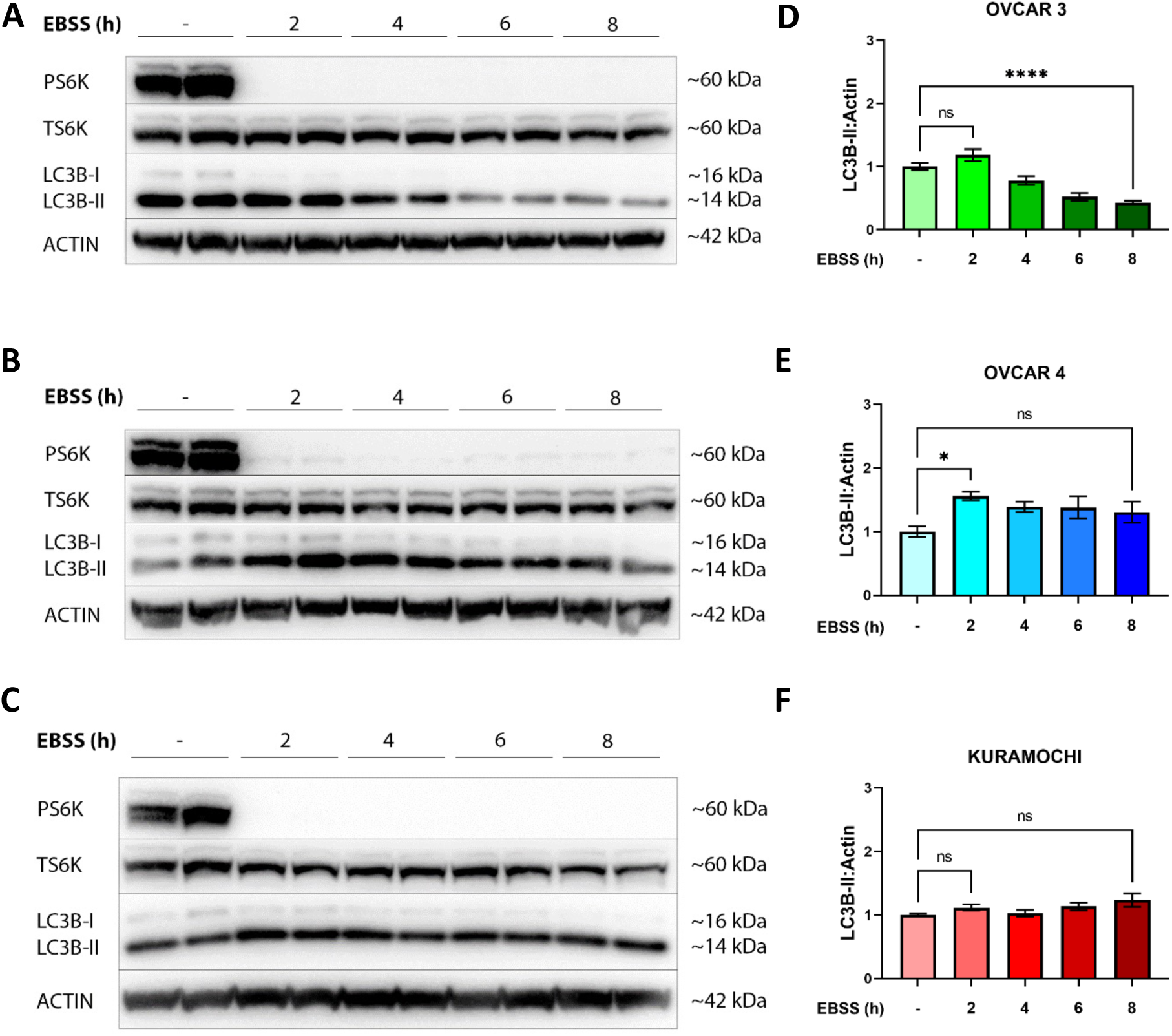
Activation of autophagy by amino acid deprivation is dysregulated in OVCAR 4 and Kuramochi cells. (**A**) OVCAR 3, (**B**) OVCAR 4 or (**C**) Kuramochi cells were incubated with EBSS for 0, 2, 4, 6 and 8 h. The phosphorylation of S6K1 at pThr389 was used to confirm the inhibition of mTORC1 signalling. β-actin was used as a loading control. (**D-F**) Autophagy flux was quantified by densitometry of LC3B-II normalised to β-actin. Densitometric analysis was pooled from three independent experiments and error bars = ±S.E.M. * p< 0.05, **** p< 0.0001 by one-way ANOVA test.

### Elevated basal autophagy in OVCAR 3 and Kuramochi, but not OVCAR 4 cells

Accumulation of autophagosomes can occur following either induction or inhibition of autophagy (17, 33) and this was resolved by measuring autophagy flux using Bafilomycin A1 (BafA1), for the final h of each EBSS timepoint (Fig. 3). OVCAR 3 and Kuramochi cells, but not OVCAR 4 cells, showed statistically significant accumulation of LC3B-II in nutrient-replete conditions, indicating a high basal autophagy activity (Fig. 3d, f).

**Figure 3.**
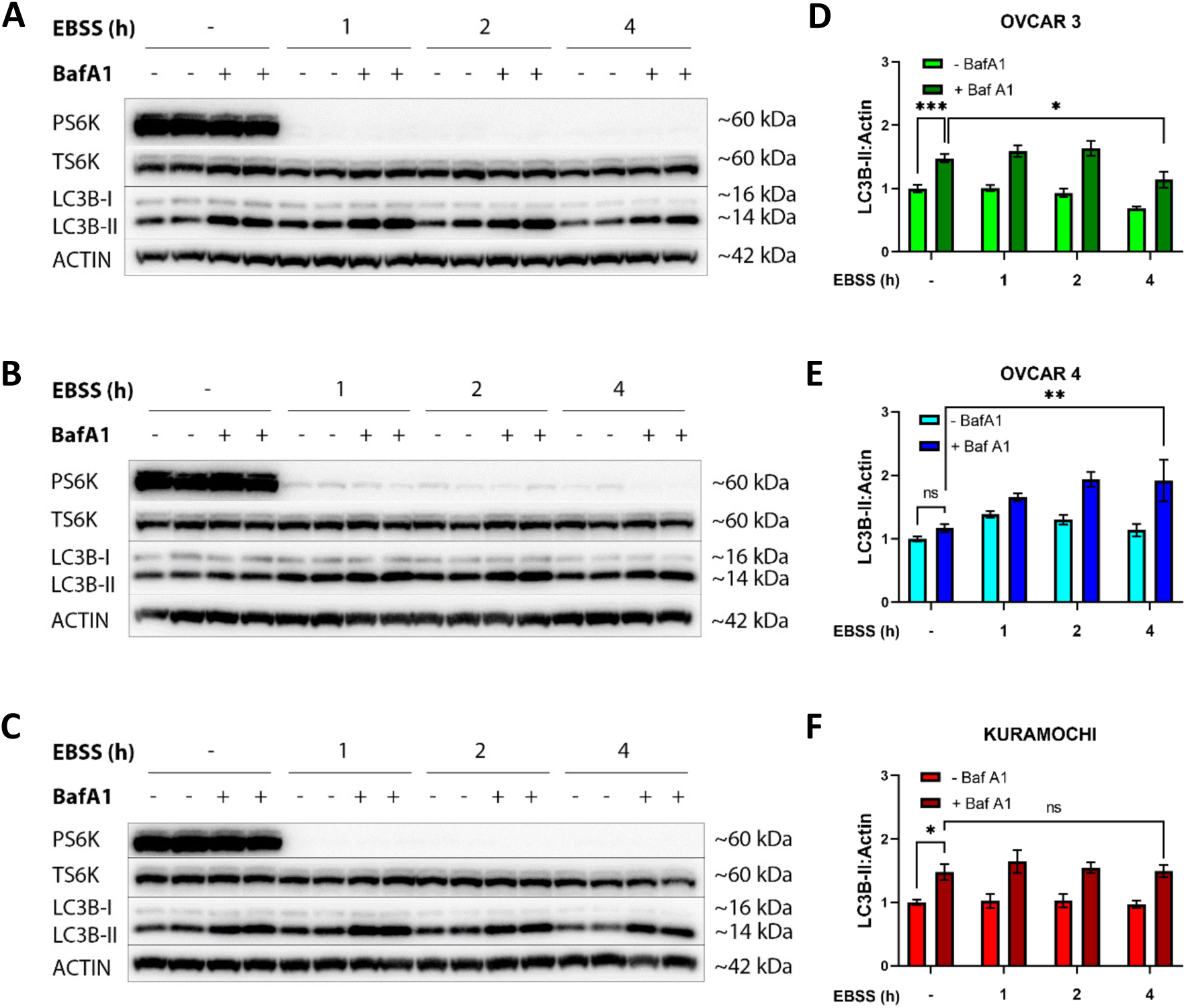
High basal autophagy is detectable in OVCAR3 and Kuramochi, but not OVCAR4 cells. (**A**) OVCAR 3, (**B**) OVCAR 4 or (**C**) Kuramochi cells were incubated with EBSS for 0, 1, 2 or 4 h before the addition of BafA1 (100 nM) for the final hour. The phosphorylation of S6K1 (pThr389) was used to confirm the inhibition of mTORC1 signalling. β-actin was used as a loading control. (**D-F**) Autophagy flux was quantified by densitometry of LC3B-II normalised to β-actin. Densitometric analysis was pooled from three independent experiments and error bars = ±S.E.M. * p< 0.05, ** p< 0.01, *** p< 0.001 by two-way ANOVA test.

Following amino acid withdrawal, OVCAR 3 cells showed a robust increase in LC3B-II accumulation after 1 and 2 h EBSS treatment although this decreased by 4 h, indicating time-dependent attenuation of autophagy (Fig. 3a, d). Conversely, OVCAR 4 cells displayed a much slower accumulation in LC3B-II up to 4 h, and LC3B-II levels remained high in the presence of BafA1, indicating that autophagy activity remained elevated even at 4 h (Fig. 3b, e). Finally, in Kuramochi cells, LC3B-II accumulation in the presence of BafA1 did not increase over time, suggesting that Kuramochi possessed basal but little activatable autophagy, at least in response to amino acid deprivation (Fig. 3c, f).

### Basal autophagy in OVCAR 3 and Kuramochi cells is RAS-dependent

Both Kuramochi and OVCAR 3 cells possess high KRAS mRNA expression (Fig. 1c) correlating with elevated basal autophagy (Fig. 3). We hypothesised that this may be another example of RAS-driven autophagy addiction in cancer (17–19). To investigate this, cells were treated with or without a MEK inhibitor to inhibit RAS signalling and autophagy flux was investigated as before, using BafA1 (Fig. 4). Following inhibition of MEK, ERK1/2 phosphorylation was decreased (Fig. 4b, d, f), as was LC3B-II accumulation in Kuramochi and OVCAR 3 cells (Fig. 4a, c), but not in OVCAR 4 cells (Fig. 4e). This data establishes that RAS-driven autophagy occurs in Kuramochi and OVCAR 3 cells.

**Figure 4.**
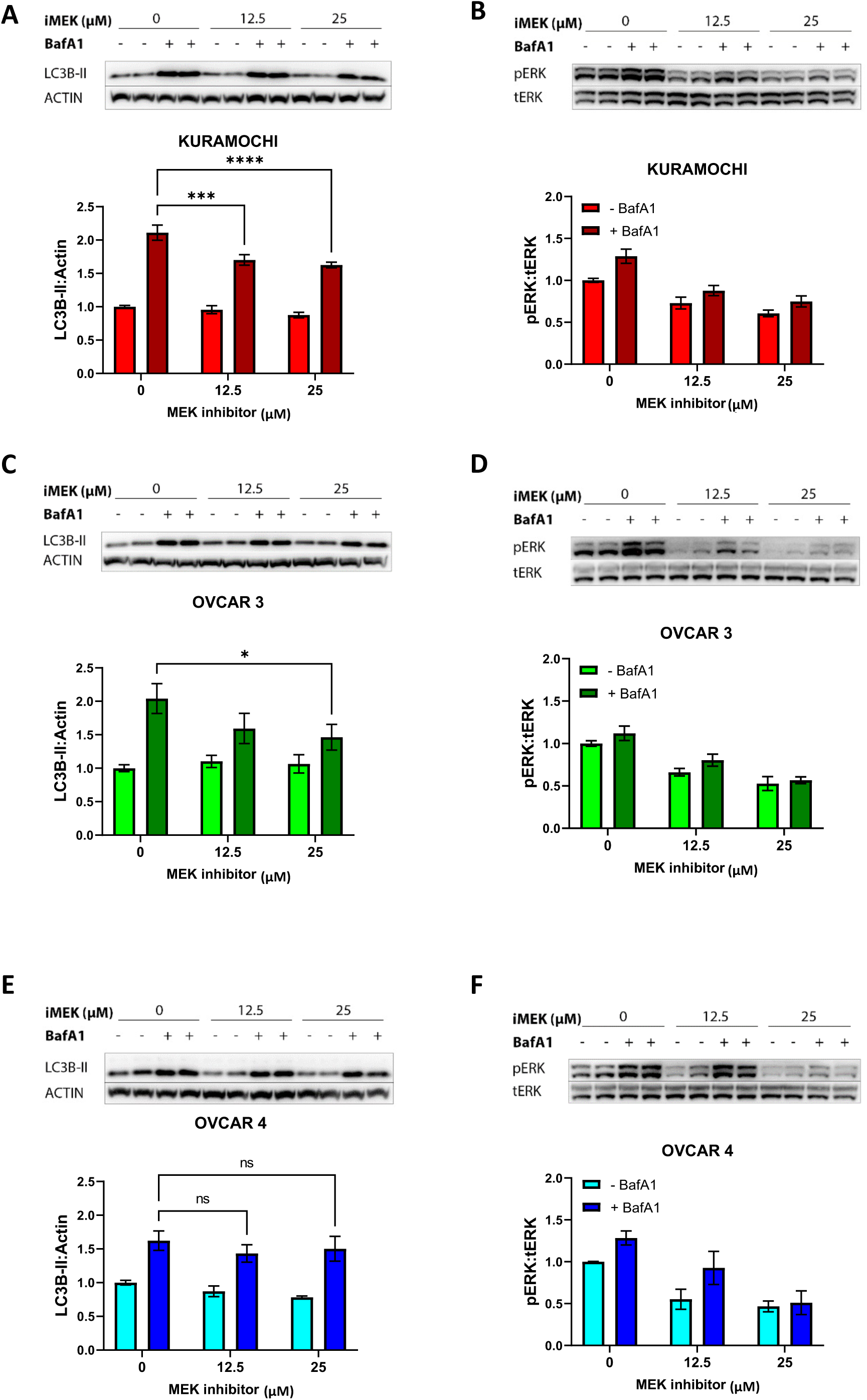
Inhibition of the Ras-pathway attenuates high basal autophagy in Kuramochi and OVCAR 3 cells. (**A, B**) Kuramochi, (**C, D**) OVCAR 3 and (**E, F**) OVCAR 4 cells were treated with MEK inhibitor, PD98059 (iMEK; 12.5 or 25 µM) for 24 h in the presence or absence of BafA1 (100 nM) for the last hour. (**A, C, E**) LC3B was probed as an autophagy marker, and β-actin was used as a loading control; Autophagy flux was quantified by densitometry of LC3B-II normalized to β-actin. (**B, D, F**) The phosphorylation of ERK1-2 at pThr202-Tyr204 was used to confirm RAS pathway inhibition and was quantified by densitometry of phosphorylated ERK normalised to total ERK expression. Densitometric analysis was pooled from a minimum of three independent experiments and error bars = ±S.E.M. * p< 0.05, *** p< 0.001, **** p< 0.0001 by two-way ANOVA test.

### PTP4A3 overexpression in OVCAR 3 cells does not promote autophagy

OVCAR 3 cells express negligible PTP4A3 (Fig. 1a-c). Previously, we reported that overexpression of PTP4A3 promoted autophagy in the non-serous OC cell line, A2780 (9). To demonstrate if PTP4A3 overexpression also enhances autophagy in OVCAR 3 cells, cells were transiently transfected with wild-type PTP4A3 (pEGFP-PTP4A3) or PTP4A1 (pEGFP-PTP4A1) and an empty vector control (pEGFPC1) (Fig. 5) (9). Increases in basal autophagy flux were assessed using BafA1. However, no statistically significant increase in LC3B-II levels was detected in cells overexpressing either PTP4A1 or PTP4A3 (Fig. 5a, b). OVCAR 3 cells overexpressing PTP4A3 were also incubated in EBSS for 2 h with or without BafA1, but again, LC3B-II levels were no different from either PTP4A1 or empty vector controls (Fig. 5c, d), revealing that PTP4A3 overexpression did not promote autophagy in OVCAR 3 HGSOC cells.

**Figure 5.**
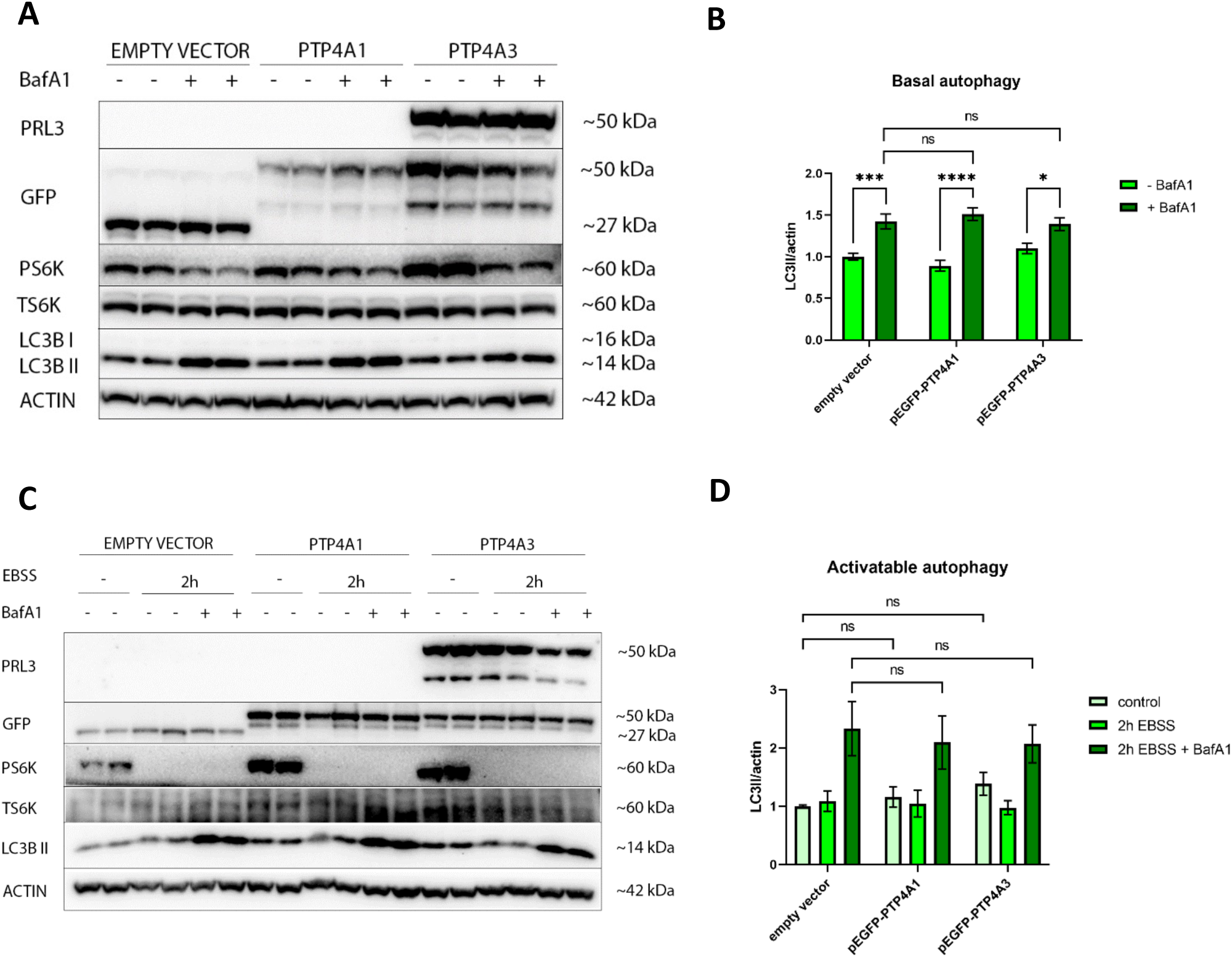
PTP4A3 overexpression in OVCAR 3 cells does not affect either basal or activatable autophagy. OVCAR 3 cells were transiently transfected with either pEGFP (empty vector), pEGFP-PTP4A1 or pEGFP-PTP4A3. (**A**) Cells were incubated with or without 100 nM BafA1 for 1 h. The phosphorylation of S6K1 (pThr389) was used to confirm the inhibition of mTORC1 signalling and β- actin was used as a loading control, while GFP was probed as a transfection efficiency control. (**B**) Basal autophagy activity was quantified by densitometry of LC3B-II and normalised to β-actin. (**C**) Alternatively, cells were incubated with or without EBSS for 2 h in the presence or absence of BafA1 (100 nM) for the last hour. (**D**) Activatable autophagy was quantified by densitometry of LC3B-II normalised to β-actin expression. Data was pooled from three independent experiments and error bars = ±S.E.M. * p< 0.05, *** p< 0.001, **** p< 0.0001 by two-way ANOVA test.

### Silencing PTP4A3 expression attenuated autophagy in OVCAR 4 cells

Since inducible and basal autophagy (Figs. 2 and 3) were dysregulated in OVCAR 4 and Kuramochi cells, the impact of PTP4A3 gene silencing on autophagy was assessed using lentivirus-mediated shRNA in Kuramochi and OVCAR 4 cells (Fig. 6a, b). In Kuramochi cells, increased LC3B-II detection was observed following BafA1 treatment under nutrient-replete conditions in both scrambled control and shPTP4A3 cells (Fig. 6c-f), indicative of high basal autophagy activity. Following amino acid withdrawal combined with BafA1, both control and shPTP4A3 treated cells showed a significant increase in LC3B-II accumulation after 1 and 2 h of EBSS treatment (Fig. 6c, d). LC3B-II accumulation was reduced after 4 h in scrambled and shPTP4A3-treated cells but was not statistically significant.

**Figure 6.**
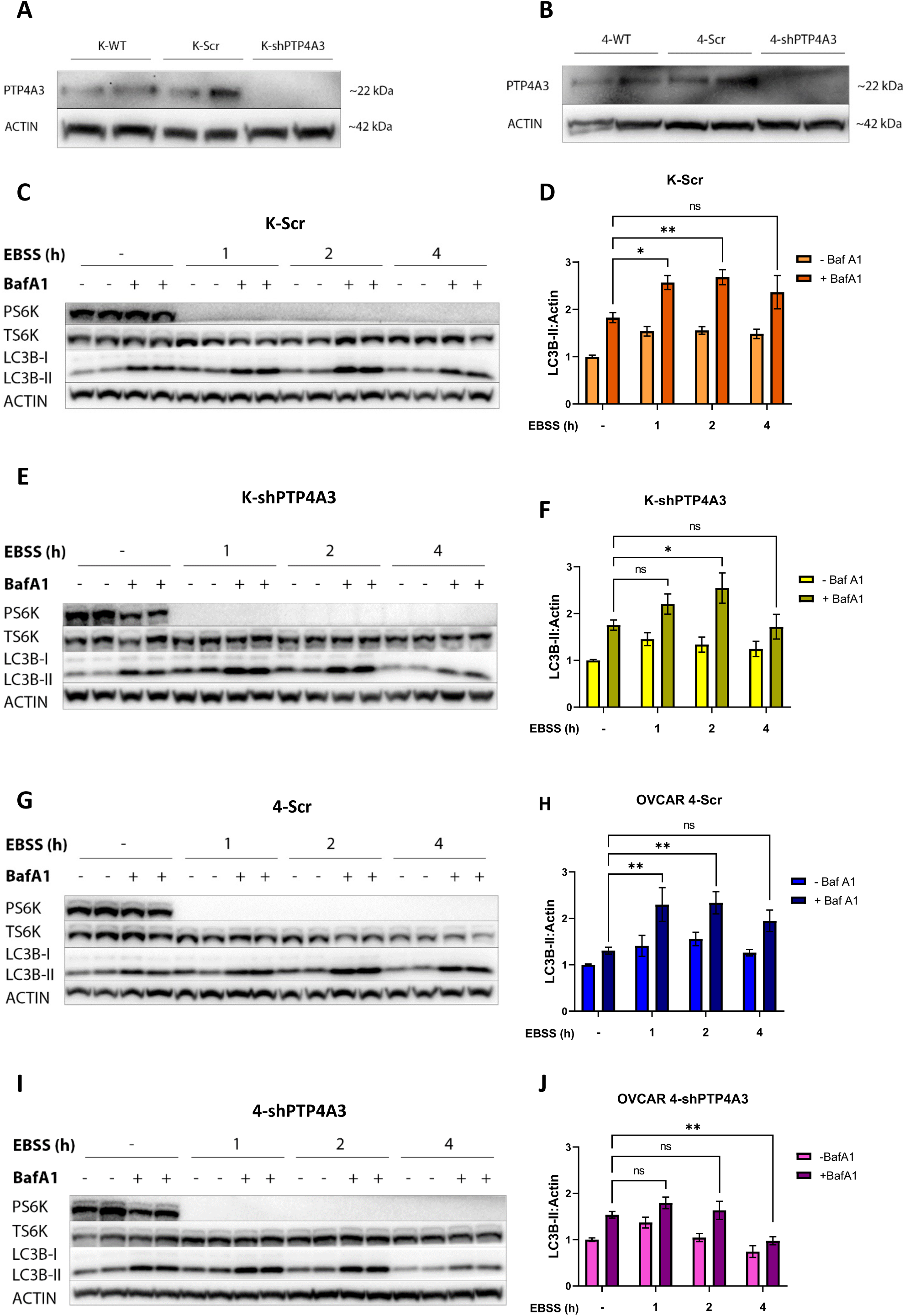
Silencing of PTP4A3 expression attenuated activatable autophagy in OVCAR 4 but not Kuramochi cells. Kuramochi (**A**) and OVCAR 4 (**B**) cells were infected with lentiviral particles harbouring either scrambled control shRNA (Scr) or PTP4A3-targeting shRNA (shPTP4A3). The silencing of PTP4A3 expression was quantified by Western immunoblotting. (**C, D**) Kuramochi-Scr (K- Scr), (**E, F**) Kuramochi-shPTP4A3 (K-shPTP4A3), (**G, H**) OVCAR 4-Scr (4-Scr) and (**I, J**) OVCAR 4-shPTP4A3 (4-shPTP4A3) cells were incubated with EBSS for 0, 1, 2 or 4 h and BafA1 (100 nM) was added for the final hour. (**C, E, G, I**) The phosphorylation of S6K1 (pThr389) was assessed by western blot to confirm inhibition of mTORC1 signalling. β-actin was used as a loading control. (**D, F, H, J**) Activation of autophagy was quantified by densitometry of LC3B-II normalised to β-actin. Data was pooled from at least three independent experiments and error bars = ±S.E.M. * p< 0.05, ** p< 0.01 by two-way ANOVA test.

In OVCAR 4 scrambled control cells, inducible autophagy was statistically significant LC3B-II accumulation following 1 and 2 h of amino acid withdrawal (Fig. 6g, h). However, in contrast, shPTP4A3 OVCAR 4 cells amino acid withdrawal did not induce autophagy, but instead, there was a significant amount of LC3B-II turnover at 4 h that did not occur in scrambled control cells (Fig. 6i, j).

### Differential PI3K and RAS signalling in Kuramochi and OVCAR 4 cells revealed by pan-inhibition of PTP4A1-3

Studies of PTP4A1 and PTP4A2 have demonstrated that those phosphatases may have overlapping functions with PTP4A3. All three PTP4A phosphatases involved in regulating cell proliferation, survival, migration, and adhesion through p53, Rho-family GTPase, PI3K, JAK-STAT, and RAS pathways, while high expression of each PTP4A is linked to several cancer types (5, 34, 35). Recently, JMS-053 (iPRL) has been identified as a reversible and non-competitive pan-PTP4A inhibitor that suppresses the migration and viability of OC cells while also preventing the *in vivo* growth of OC xenografts (36, 37).

PTP4A3 is the most frequently overexpressed PTP4A phosphatase in different cancer types, although some cancers also co-express PTP4A1 and/or PTP4A2 (5, 34). This suggests that compensatory mechanisms may arise when therapeutically targeting a single PTP4A species in isolation. To understand whether inhibition of all PTP4A proteins altered oncogenic signalling, we analysed the effect of iPRL on cell signalling in the Kuramochi and OVCAR 4 cell lines, with and without PTP4A3 knockdown.

Incubation of Kuramochi cells with ≥ 5 µM iPRL activated PI3K and RAS signalling, detected by increased AKT and ERK1/2 phosphorylation (Fig. 7a). In contrast, iPRL inhibited PI3K and RAS signalling in OVCAR 4 cells (Fig. 7b). Strikingly, ≥ 5 µM iPRL inhibited S6K1 phosphorylation in WT, scrambled control and shPTP4A3 knockdowns of both cell lines (Fig. 7a, b), which was associated with decreased ULK1 phosphorylation at Ser 757, the mTORC1 inhibitory site, suggesting that targeting pan-PTP4As inhibited mTORC1 at a point distal to AKT signalling.

**Figure 7.**
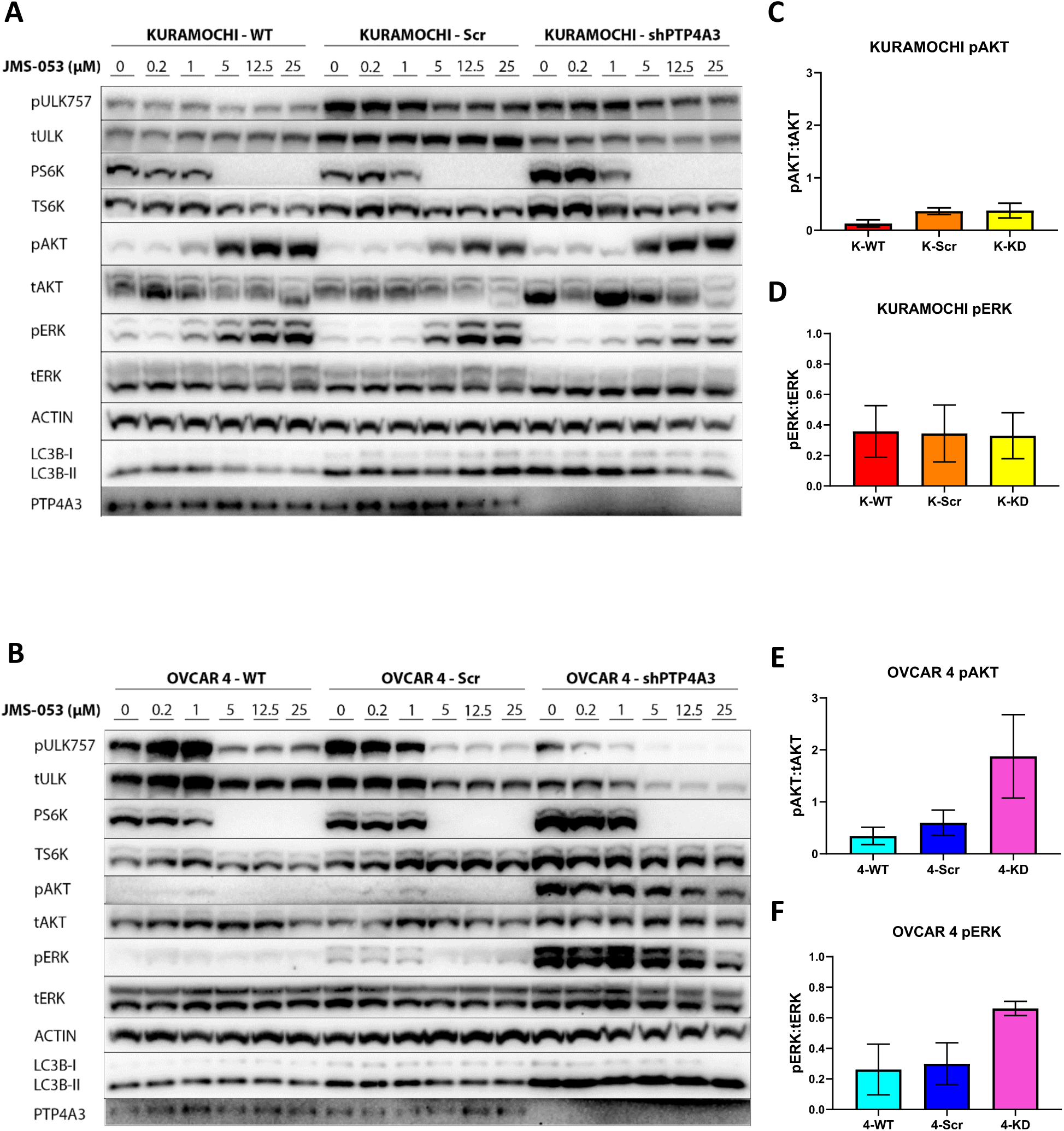
Inhibition of PTP4A1-3 with JMS-053 revealed differential PI3K and RAS signalling in Kuramochi and OVCAR 4 cells. (**A**) Kuramochi-WT, Scr and shPTP4A3 and (**B**) OVCAR 4-WT, Scr and shPTP4A3 cells were treated with increasing concentrations of JMS-053 (iPRL; 0 - 25 µM) for 2 h. Cell extracts were analysed for phosphorylation of ULK1 (pSer757), S6K1 (pThr389), AKT/PKB (pSer473) and ERK1-2 (pThr202-Tyr204). LC3B was used as a marker of autophagy activity, β-actin was used as a loading control, and PTP4A3 knockdown was also confirmed. (**C-F**) The basal phosphorylation of AKT and ERK in WT, Scr and shPTP4A3 cells (no JMS-053 treatment) in (**C, D**) Kuramochi and (**E, F**) OVCAR 4 cells was determined. Data was pooled from three independent experiments and error bars = ±S.E.M.

Distinct differences in PI3K and RAS signalling were observed in OVCAR 4 cells following PTP4A3 silencing (Fig. 7b). Quantification of AKT and ERK1/2 phosphorylation revealed no change in Kuramochi shPTP4A3 cells compared to control cells (Fig. 7c, d). Conversely, silencing PTP4A3 in OVCAR 4 cells elevated both AKT (Ser473) and ERK1/2 phosphorylation, although these were not statistically significant (Fig. 7e, f).

### PTP4A3 knockdown sensitises K-Ras mutant HGSOC to 5-fluorouracil

Most women with advanced OC initially respond favourably to first-line treatments, carboplatin and paclitaxel. However, patients develop chemoresistance leading to treatment failure in 80-90% of cases, with a 5-year survival rate of < 40 % for women with HGSOC (38–41). OC heterogeneity represents a major challenge for the improvement of cure rates for this disease (38, 40). The HGSOC cell lines used in this study display different levels of PTP4A3 and KRAS expression. To translate our results towards improved therapeutic strategies, we assessed whether PTP4A3 knockdown sensitised HGSOC cells towards pan-PTP4A inhibition (iPRL) or to the clinically relevant chemotherapeutic agents, fluorouracil (5FU), cisplatin (CDDP), and paclitaxel (PTX).

We employed time-lapse cell culture imaging and analysis to quantify cell confluency and incorporation of propidium iodide (PI), as measures of reduced cell proliferation and cell death, respectively. We found that Kuramochi cells may be more dependent on PTP4A3 activity given their higher PTP4A3 expression which resulted in greater sensitivity towards iPRL, compared to OVCAR 4 and OVCAR 3 (Supplementary Fig. S1) while PTP4A3 silencing in both Kuramochi and OVCAR 4 cell lines also appeared to sensitise these cells to iPRL (Supplementary Figures S2 and S3). Moreover, the absence of PTP4A3 expression was associated with greater sensitivity to chemotherapeutic drugs. OVCAR 3 cells were more sensitive to CDDP and PTX than either WT Kuramochi or OVCAR 4 cells (Supplementary Figures S7 and S10). The shPTP4A3 Kuramochi cell line was more sensitive to 5-FU, CDDP and PTX than their control counterparts (Supplementary Figures S5, S8 and S11), while shPTP4A3 OVCAR 4 cells were more sensitive towards 5-FU and PTX (Supplementary Figures S6 and S12).

Compared to scrambled control, shPTP4A3 Kuramochi cells treated with 5 µM iPRL had significantly lower cell confluency at 48 h (Fig. 8a). Similarly, silencing PTP4A3 in Kuramochi cells sensitised them towards 100 µM 5-FU (Fig. 8c). We also observed a trend of increased sensitivity to 10 µM CDDP (Fig. 8e) and 10 nM PTX (Fig. 8g) in Kuramochi PTP4A3 knockdown cells. However, silencing PTP4A3 in OVCAR 4 cells did not alter drug sensitivity compared to scrambled controls (Fig. 8b, d, f, h). Nonetheless, the calculated IC_50_ values for each drug revealed increased sensitivity towards 5-FU and CDDP following PTP4A3 silencing in Kuramochi and OVCAR 4 cells (Fig. 8i and Supplementary Fig. S13). We also noted that iPRL-treated shPTP4A3 Kuramochi cells had a 4-fold lower IC_50_ compared to scrambled control (Fig. 8i and Supplementary Fig. S13), suggesting that compensatory mechanisms occur between PTP4A proteins.

**Figure 8.**
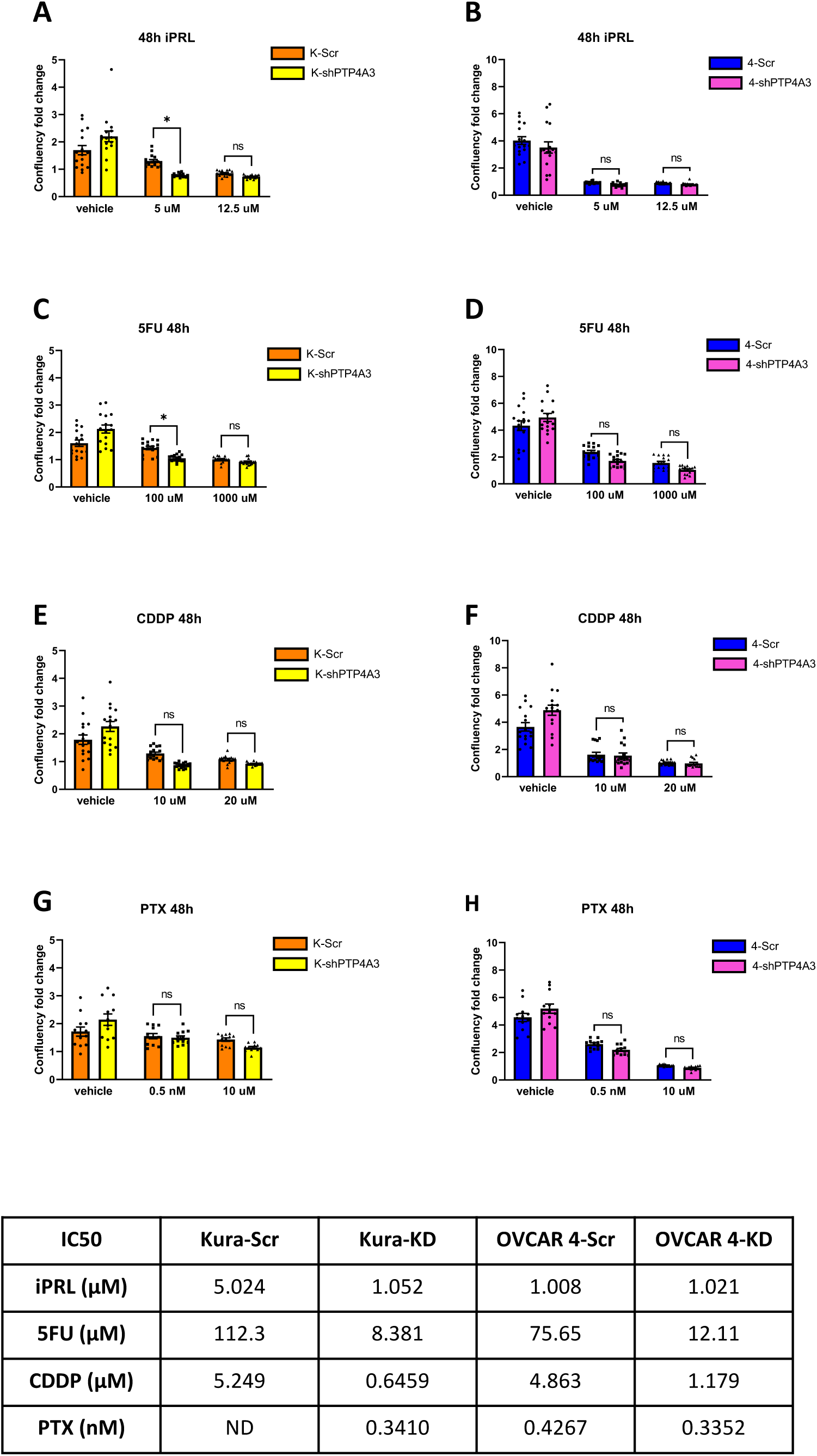
Silencing of PTP4A3 sensitises K-Ras mutant, but not K-Ras wildtype HGSOC to iPRL and 5FU. K-Scr, K-shPTP4A3, OVCAR 4-Scr and OVCAR 4-shPTP4A3 cells were treated with; (**A, B**) JMS-053 (iPRL), (**C, D**) 5- fluorouracil (5FU), (**E, F**) cisplatin (CDDP) or (**G, H**) paclitaxel (PTX) for 72 h in an Incucyte S3 system. (**A-H**) Cell confluency at 48 h relative to T_0_. Data was pooled from three (PTX) or four (iPRL, 5FU and CDDP) independent experiments and error bars = ±S.E.M., with * p< 0.05 by two-way ANOVA test. (**I**) The IC_50_ values for each drug.

## DISCUSSION

Numerous studies have reported the involvement of PTP4A3 in malignant transformation and EMT through several oncogenic pathways, including the PI3K, RAS and SRC pathways (24, 42). Further, we previously demonstrated the ability of PTP4A3 overexpression to promote autophagy in A2780 cells (9). Thus, we sought to delineate the molecular mechanisms by which PTP4A3 mediates cell growth, proliferation and autophagy in HGSOC cells.

Analysis of RNA and protein expression revealed PTP4A3 expression levels varies across HGSOC cell lines: Kuramochi (high), OVCAR 4 (low) and OVCAR 3 cells (negligible) (Fig. 1a, b, c). In Kuramochi cells, the previously reported KRAS amplification (28) correlated with elevated levels of ERK phosphorylation (Fig. 1c, d). Although oncogenic RAS mutations activate both PI3K and RAS signalling pathways, KRAS is more effective in inducing RAS signalling whereas HRAS more potently activates the PI3K pathway (43, 44). Therefore, for Kuramochi cells, high KRAS in combination with high PTP4A3 expression may result in an over-activation of the RAS pathway, which resulted in the observed low PI3K signalling activity we detected, due to known pathway crosstalk (45). Conversely, PTP4A3 expression and the absence of KRAS amplification in OVCAR 4 cells may be the reason why this cell line showed high PI3K and RAS activity, although RAS activity was still less than in Kuramochi cells. OVCAR 3 cells did not express PTP4A3, while their elevated KRAS mRNA expression was still below those of Kuramochi cells (Figure 1c), these two features may explain, in part, the low RAS and PI3K pathway activity, compared to Kuramochi and OVCAR 4 cells (Figure 1d).

Long-term amino acid deprivation combined with autophagy flux analysis using BafA1, revealed that OVCAR 3 cells possess high basal and activatable autophagy levels, whereas OVCAR 4 cells had both low basal and low activatable autophagy profiles, and Kuramochi cells displayed high basal but no activatable autophagy activity (Figs. 2 and 3). Elevated basal and minimal activatable autophagy is a common phenotype of KRAS-driven, autophagy-addicted cancers (16–19). The RAS pathway controls proliferation, differentiation, and growth (44, 46), and mutant RAS promotes basal autophagy and is thought to sustain tumour-cell mitochondrial metabolism mechanistically through a mTORC1- independent autophagy activity. Therefore, mTORC1 inhibition does not further increase autophagy in RAS-driven ‘autophagy addicted’ tumour cells (17–19). However, the level of ERK1/2 phosphorylation (Fig. 1d) observed did not correlate with known high KRAS expression and elevated basal autophagy in OVCAR 3 cells. Despite this, the MEK inhibitor was able to reduce basal autophagy in both OVCAR 3 and Kuramochi cells but not in OVCAR 4 cells, suggesting that basal autophagy in Kuramochi and OVCAR 3 cells was KRAS-driven (Fig. 4).

Further investigation of the role of PTP4A3 in regulating autophagy in HGSOC revealed that transient transfection of PTP4A3 into OVCAR 3 cells did not affect basal (Fig. 5a, b) or a.a. deprivation-induced activatable autophagy (Fig. 5c, d), which contrast with our previous findings in A2780 cells (9). However, the high KRAS expression and RAS-dependent autophagy in OVCAR 3 cells may mask any autophagy-promoting effects of PTP4A3 overexpression in this cell line. While silencing of PTP4A3 in Kuramochi cells did not alter autophagy (Fig. 6c-f), knockdown of PTP4A3 in OVCAR 4 cells led to decreased activatable autophagy (Fig. 6g-j) suggesting that endogenous PTP4A3 expression in OVCAR 4 cells is necessary for autophagy induction, at least for amino acid withdrawal.

We addressed potential compensatory mechanisms between PTP4A1-3 by employing JMS-053, a pan-PRL inhibitor (iPRL). Treatment of Kuramochi cells with ≥ 5 μM iPRL increased PI3K and RAS pathway signalling (Fig. 7a). Conversely, in OVCAR 4 cells iPRL decreased PI3K or RAS pathway signalling, at the same concentration (Fig. 7b). While few small molecule inhibitors can claim absolute specificity, we were still surprised to observe that iPRL treatment led to attenuated S6K1 and ULK1 phosphorylation in all cell lines tested, revealing a previously unreported inhibition of mTORC1. Moreover, the PTP4A3 knockdown cell lines showed a similar response to iPRL compared to control cells. These cells also express PTP4A1 and/or PTP4A2, in agreement with RNAseq analysis (Fig. 1c) and that all three phosphatases participate in the regulation of RAS and PI3K signalling, and potentially also independently, mTORC1. Thus, targeting only PTP4A3 only, via shRNA silencing resulted in the upregulation of compensatory mechanisms (Fig. 7e, f). Conversely, since high KRAS expression appears to alter these signalling pathways in Kuramochi, silencing of PTP4A3 did not elicit these effects in this HGSOC line (Figure 7c, d). PTP4A1-3 inhibition in Kuramochi was only efficient at inhibiting mTORC1 activity, suggesting that PTP4As function at different points along the PI3K and RAS pathways, including proximal to mTORC1, while high KRAS expression did not impact mTORC1 activity.

Finally, we considered the reported roles of PTP4A3 in tumour recurrence, and chemotherapy resistance, where it is upregulated following exposure to chemotherapeutic drugs (47, 48). Assays of cell proliferation and death revealed that the highest expression of PTP4A3, found in Kuramochi cells, also correlated with the greatest sensitivity to iPRL (Supplementary Fig. S1). Additionally, the absence of PTP4A3 in OVCAR 3 cells corresponded to increased cytotoxicity from CDDP (Supplementary Fig. S7) and PTX (Supplementary Fig. S10) when compared to Kuramochi and OVCAR 4 cells. However, while the shPTP4A3 Kuramochi cell line was more sensitive to 5-FU, CDDP and PTX (Supplementary Figs. S5, S8 and S11), and shPTP4A3 OVCAR 4 cells were more sensitive to 5- FU and PTX (Supplementary Figs. S6 and S12) compared to scramble controls, only shPTP4A3 Kuramochi cells showed a statistically significant sensitisation to iPRL and 5FU (Fig. 8a, c). Nevertheless, the IC_50_ values for 5FU and CDDP were ≥4 times lower in shPTP4A3 OVCAR 4 and shPTP4A3 Kuramochi cells and also for iPRL in shPTP4A3 Kuramochi cells. These results suggest that targeting HGSOC that have PTP4A3 expression sensitises them to DNA damaging chemotherapeutics and this approach is also more effective in KRAS mutant HGSOC cells.

This study has demonstrated that compensatory mechanisms from PTP4A1 and PTP4A2 can arise when specifically targeting PTP4A3 in HGSOC and that pan-PTP4A inhibition can overcome those effects. Moreover, KRAS-driven tumours may compensate for the lack of PTP4A1-3 activity by increasing RAS and PI3K signalling, coupled with autophagy addiction. Importantly however, a combinatorial approach of targeting PTP4A3 alongside current generation, clinically used chemotherapeutic drugs (5-fluorouracil, cisplatin and paclitaxel) could be an effective strategy for treating KRAS mutant, HGSOC cancers.

## ACKNOWLEDGEMENTS

This work was funded by a Scholarship from the Irish Research Council (GOIPG/2018/2724).

## AUTHOR CONTRIBUTIONS

ALG contributed to the conceptualization of experiments, the generation of in vitro data and the analysis and interpretation of results. ALG also wrote the manuscript. DJ was responsible for generating the normalized RNA-seq data and its analysis. EC was responsible for the analysis of experimental data, interpreting results, and writing and providing feedback on the manuscript. JTM was responsible for conceptualizing the experimental approach, generation, and analysis of experimental data, interpreting the results, and writing and providing feedback on the manuscript.

## DATA AVAILABILITY STATEMENT

All relevant materials and reagents are freely available upon request from the corresponding author, as are all raw data to any researcher wishing to use them for non-commercial purposes.

