## Supplemental data for "Targeting of PTP4A3 overexpression sensitises HGSOC cells towards chemotherapeutic drugs"

#### SUPPLEMENTARY FIGURES

**Supplementary figure 1. IC50 curves for JMS-053, 5FU, CDDP and PTX.** Confluency data at 48 h from the previous experiment were analysed to obtain the IC50s for each drug in Kuramochi and OVCAR 4 comparing shPTP4A3 cell lines against their scrambled control counterparts. Data was pooled from 3 (PTX) or 4 (iPRL, 5FU and CDDP) independent experiments, error bars = ±S.E.M.

**Supplementary figure 2. Kuramochi cells show higher sensitivity to the pan-PTP4A/PRL inhibitor (iPRL) than OVCAR 3 and OVCAR 4 cells.** Kuramochi, OVCAR 4 and OVCAR 3 cells were treated with increasing iPRL (JMS-053) concentrations (0 to 25 µM) in the presence of 2.5 µg/ml Propidium Iodide (PI) and incubated in the Incucyte for 72 hrs. Phase-contrast and red fluorescence (PI) images were taken every 3 hrs. A) Cell confluence plotted over time relative to initial (0 hr) confluence (set to 100) (left), and the ratio of red fluorescence (PI-positive) to total cell confluence plotted over time (right). Comparative analysis of the 3 HGSOC cell lines treated with 0 µM, 5 µM and 12.5 µM iPRL for B) confluency over 72 hrs and C) 24 hrs for PI/confluency (relative to control (0 hr)). Data was pooled from 3 (Kuramochi) or 4 (OVCAR 3 and OVCAR 4) independent experiments, error bars = ±S.E.M. D) Overlayed phase-contrast and fluorescence images (10x magnification) at 0 and 24 hrs treatment with 0 µM (0.2% DMSO), 5 µM or 12.5 µM iPRL.

**Supplementary figure 3. Kuramochi-KD (K-KD) cells show higher sensitivity to PRL inhibitor (iPRL) than K-Scr cells, however, K-Scr shows higher resistance than K-WT cells.** K-WT, K-Scr and K-KD cells were treated with increasing iPRL (JMS-053) concentrations (0 to 25 µM) in the presence of 2.5 µg/ml Propidium Iodide (PI) and incubated in the Incucyte for 72 hrs. Phase-contrast and red fluorescence (PI) images were taken every 3 hrs. **A)** Cell confluence plotted over time relative to initial (0 hr) confluence (set to 100) (left), and the ratio of red fluorescence (PI-positive) to total cell confluence plotted over time (right). Comparative analysis of the 3 Kuramochi cell lines treated with 0 µM, 5 µM and 12.5 µM iPRL for: **B)** confluency over 72 hrs and **C)** 24 hrs for PI/confluency (relative to control (0 hr)). Data was pooled from 3 (K-WT) or 4 (K-Scr and K-KD) independent experiments, error bars = ±S.E.M. **D)** Overlayed phase-contrast and fluorescence images (10x magnification) at 0 and 24 hrs treatment with 0 µM (0.2% DMSO), 5 µM or 12.5 µM iPRL.

**Supplementary figure 4. OVCAR 4-KD (4-KD) cells show higher sensitivity to the PRL inhibitor (iPRL) than 4-WT and 4-Scr cells.** 4-WT, 4-Scr and 4-KD cells were treated with increasing iPRL (JMS-053) concentrations (0 to 25 µM) in the presence of 2.5 µg/ml Propidium Iodide (PI) and incubated in the Incucyte for 72 hrs. Phase-contrast and red fluorescence (PI) images were taken every 3 hrs. **A)** Cell confluence plotted over time relative to initial (0 hr) confluence (set to 100) (left), and the ratio of red fluorescence (PI-positive) to total cell confluence plotted over time (right). Comparative analysis of the 3 OVCAR 4 cell lines treated with 0 µM, 5 µM and 12.5 µM iPRL for: **B)** confluency over 72 hrs and **C)** 24 hrs for PI/confluency (relative to control (0 hr)). Data was pooled from 4 independent experiments, error bars = ±S.E.M. **D)** Overlayed phase-contrast and fluorescence images (10x magnification) at 0 and 24 hrs treatment with 0 µM (0.2% DMSO), 5 µM or 12.5 µM iPRL.

**Supplementary figure 5. Kuramochi cells show higher sensitivity to 5FU than OVCAR 3 and OVCAR 4.** Kuramochi, OVCAR 4 and OVCAR 3 cells were treated with increasing 5FU concentrations (0 to 1 mM) in the presence of 2.5 µg/ml Propidium Iodide (PI) and incubated in the Incucyte for 72 hrs. Phase-contrast and red fluorescence (PI) images were taken every 3 hrs. **A)** Cell confluence plotted over time relative to initial (0 hr) confluence (set to 100) (left), and the ratio of red fluorescence (PI-positive) to total cell confluence plotted over time (right). Comparative analysis of the 3 HGSOC cell lines treated with 0 µM, 100 µM and 1000 µM 5FU for: **B)** confluency over 72 hrs and **C)** 24 hrs for PI/confluency (relative to control (0 hr)). Data was pooled from 4 independent experiments, error bars = ±S.E.M. **D)** Overlayed phase-contrast and fluorescence images (10x magnification) at 0 and 24 hrs treatment with 0 µM (0.5% DMSO), 100 µM and 1 mM 5FU.

**Supplementary figure 6. Kuramochi-KD (K-KD) cells show higher sensitivity to 5FU than K-Scr, however, K-Scr shows higher resistance than K-WT.** K-WT, K-Scr and K-KD cells were treated with increasing 5FU concentrations (0 to 1 mM) in the presence of 2.5 µg/ml Propidium Iodide (PI) and incubated in the Incucyte for 72 hrs. Phase-contrast and red fluorescence (PI) images were taken every 3 hrs. **A)** Cell confluence plotted over time relative to initial (0 hr) confluence (set to 100) (left), and the ratio of red fluorescence (PI-positive) to total cell confluence plotted over time (right). Comparative analysis of the 3 Kuramochi cell lines treated with 0 µM, 100 µM and 1000 µM 5FU for: **B)** confluency over 72 hrs and **C)** 24 hrs for PI/confluency (relative to control (0 hr)). Data was pooled from 4 independent experiments, error bars = ±S.E.M. **D)** Overlayed phase-contrast and fluorescence images (10x magnification) at 0 and 24 hrs treatment with 0 µM (0.5% DMSO), 100 µM and 1 mM 5FU.

**Supplementary figure 7. OVCAR 4-KD (4-KD) cells show higher sensitivity to 5FU than 4-WT and 4-Scr.** 4-WT, 4-Scr and 4-KD cells were treated with increasing 5FU concentrations (0 to 1 mM) in the presence of 2.5 µg/ml Propidium Iodide (PI) and incubated in the Incucyte for 72 hrs. Phase-contrast and red fluorescence (PI) images were taken every 3 hrs. **A)** Cell confluence plotted over time relative to initial (0 hr) confluence (set to 100) (left), and the ratio of red fluorescence (PI-positive) to total cell confluence plotted over time (right). Comparative analysis of the 3 OVCAR 4 cell lines treated with 0 µM, 100 µM and 1000 µM 5FU for: **B)** confluency over 72 hrs and **C)** 24 hrs for PI/confluency (relative to control (0 hr)). Data was pooled from 4 independent experiments, error bars = ±S.E.M. **D)** Overlayed phase-contrast and fluorescence images (10x magnification) at 0 and 24 hrs treatment with 0 µM (0.5% DMSO), 100 µM and 1 mM 5FU.

**Supplementary figure 8. OVCAR 3 cells show higher sensitivity to cisplatin (CDDP) than OVCAR 4 and Kuramochi.** Kuramochi, OVCAR 4 and OVCAR 3 cells were treated with increasing CDDP concentrations (0 to 40 µM) in the presence of 2.5 µg/ml Propidium Iodide (PI) and incubated in the Incucyte for 72 hrs. Phase-contrast and red fluorescence (PI) images were taken every 3 hrs. **A)** Cell confluence plotted over time relative to initial (0 hr) confluence (set to 100) (left), and the ratio of red fluorescence (PI-positive) to total cell confluence plotted over time (right). Comparative analysis of the 3 HGSOC cell lines treated with 0 µM, 10 µM and 20 µM CDDP for: **B)** confluency over 72 hrs and **C)** 24 hrs for PI/confluency (relative to control (0 hr)). Data was pooled from 4 independent experiments, error bars = ±S.E.M. **D)** Overlayed phase-contrast and fluorescence images (10x magnification) at 0 and 24 hrs treatment with 0 µM (5% of 0.9% NaCl), 10 µM and 20 µM CDDP.

**Supplementary figure 9. Kuramochi-KD (K-KD) cells show higher sensitivity to cisplatin (CDDP) than K-WT and K-Scr.** K-WT, K-Scr and K-KD cells were treated with increasing CDDP concentrations (0 to 40 µM) in the presence of 2.5 µg/ml Propidium Iodide (PI) and incubated in the Incucyte for 72 hrs. Phase-contrast and red fluorescence (PI) images were taken every 3 hrs. **A)** Cell confluence plotted over time relative to initial (0 hr) confluence (set to 100) (left), and the ratio of red fluorescence (PI-positive) to total cell confluence plotted over time (right). Comparative analysis of the 3 Kuramochi cell lines treated with 0 µM, 10 µM and 20 µM CDDP for: **B)** confluency over 72 hrs and **C)** 24 hrs for PI/confluency (relative to control (0 hr)). Data was pooled from 4 independent experiments, error bars = ±S.E.M. **D)** Overlayed phase-contrast and fluorescence images (10x magnification) at 0 and 24 hrs treatment with 0 µM (5% of 0.9% NaCl), 10 µM and 20 µM CDDP.

**Supplementary figure 10. *PTP4A3* silencing does not produce a significant effect in the response of OVCAR 4 cells to cisplatin (CDDP) treatment.** 4-WT, 4-Scr and 4-KD cells were treated with increasing CDDP concentrations (0 to 40 µM) in the presence of 2.5 µg/ml Propidium Iodide (PI) and incubated in the Incucyte for 72 hrs. Phase-contrast and red fluorescence (PI) images were taken every 3 hrs. **A)** Cell confluence plotted over time relative to initial (0 hr) confluence (set to 100) (left), and the ratio of red fluorescence (PI-positive) to total cell confluence plotted over time (right). Comparative analysis of the 3 OVCAR 4 cell lines treated with 0 µM, 10 µM and 20 µM CDDP for: **B)** confluency over 72 hrs and **C)** 24 hrs for PI/confluency (relative to control (0 hr)). Data was pooled from 4 independent experiments, error bars = ±S.E.M. **D)** Overlayed phase-contrast and fluorescence images (10x magnification) at 0 and 24 hrs treatment with 0 µM (5% of 0.9% NaCl), 10 µM and 20 µM CDDP.

**Supplementary figure 11. OVCAR 3 cells show higher sensitivity to paclitaxel (PTX) than OVCAR 4 and Kuramochi.** Kuramochi, OVCAR 4 and OVCAR 3 cells were treated with increasing PTX concentrations (0 to 100 nM) in the presence of 2.5 µg/ml Propidium Iodide (PI) and incubated in the Incucyte for 72 hrs. Phase-contrast and red fluorescence (PI) images were taken every 3 hrs. **A)** Cell confluence plotted over time relative to initial (0 hr) confluence (set to 100) (left), and the ratio of red fluorescence (PI-positive) to total cell confluence plotted over time (right). Comparative analysis of the 3 HGSOC cell lines treated with 0 nM, 0.5 nM and 10 nM PTX for: **B)** confluency over 72 hrs and **C)** 24 hrs for PI/confluency (relative to control (0 hr)). Data was pooled from 3 independent experiments, error bars = ±S.E.M. **D)** Overlayed phase-contrast and fluorescence images (10x magnification) at 0 and 24 hrs treatment with 0 nM (0.1% DMSO), 0.5 nM and 10 nM PTX.

**Supplementary figure 12. Kuramochi-KD (K-KD) cells show higher sensitivity to paclitaxel (PTX) than K-Scr, however, K-WT is the most sensitive.** K-WT, K-Scr and K-KD cells were treated with increasing PTX concentrations (0 to 100 nM) in the presence of 2.5 μg/ml Propidium Iodide (PI) and incubated in the Incucyte for 72 hrs. Phase-contrast and red fluorescence (PI) images were taken every 3 hrs. **A)** Cell confluence plotted over time relative to initial (0 hr) confluence (set to 100) (left), and the ratio of red fluorescence (PI-positive) to total cell confluence plotted over time (right). Comparative analysis of the 3 Kuramochi cell lines treated with 0 nM, 0.5 nM and 10 nM PTX for: **B)** confluency over 72 hrs and **C)** 24 hrs for PI/confluency (relative to control (0 hr)). Data was pooled from 3 independent experiments, error bars = ±S.E.M. **D)** Overlayed phase-contrast and fluorescence images (10x magnification) at 0 and 24 hrs treatment with 0 nM (0.1% DMSO), 0.5 nM and 10 nM PTX.

**Supplementary figure 13. OVCAR 4-KD (4-KD) cells show higher sensitivity to paclitaxel (PTX) than 4-WT and 4-Scr.** 4-WT, 4-Scr and 4-KD cells were treated with increasing PTX concentrations (0 to 100 nM) in the presence of 2.5 μg/ml Propidium Iodide (PI) and incubated in the Incucyte for 72 hrs. Phase-contrast and red fluorescence (PI) images were taken every 3 hrs. **A)** Cell confluence plotted over time relative to initial (0 hr) confluence (set to 100) (left), and the ratio of red fluorescence (PI-positive) to total cell confluence plotted over time (right). Comparative analysis of the 3 OVCAR 4 cell lines treated with 0 nM, 0.5 nM and 10 nM PTX for: **B)** confluency over 72 hrs and **C)** 24 hrs for PI/confluency (relative to control (0 hr)). Data was pooled from 3 independent experiments, error bars = ±S.E.M. **D)** Overlayed phase-contrast and fluorescence images (10x magnification) at 0 and 24 hrs treatment with 0 nM (0.1% DMSO), 0.5 nM and 10 nM PTX.

**Supplementary figure 14. PTP4A3 mRNA expression in OVCAR 4 and Kuramochi cells upon lentiviral mediated shRNA knockdown.** RNA from OVCAR 4 (top) and Kuramochi (bottom) WT, Scrambled and Knockdowns (1883 and 1884) cell lysates were isolated and analysed by qPCR. PTP4A3 expression levels were normalised to GAPDH and are depicted as a fold increase. Data represents three independent experiments, error bars = ± S.E.M.

### Figure S1

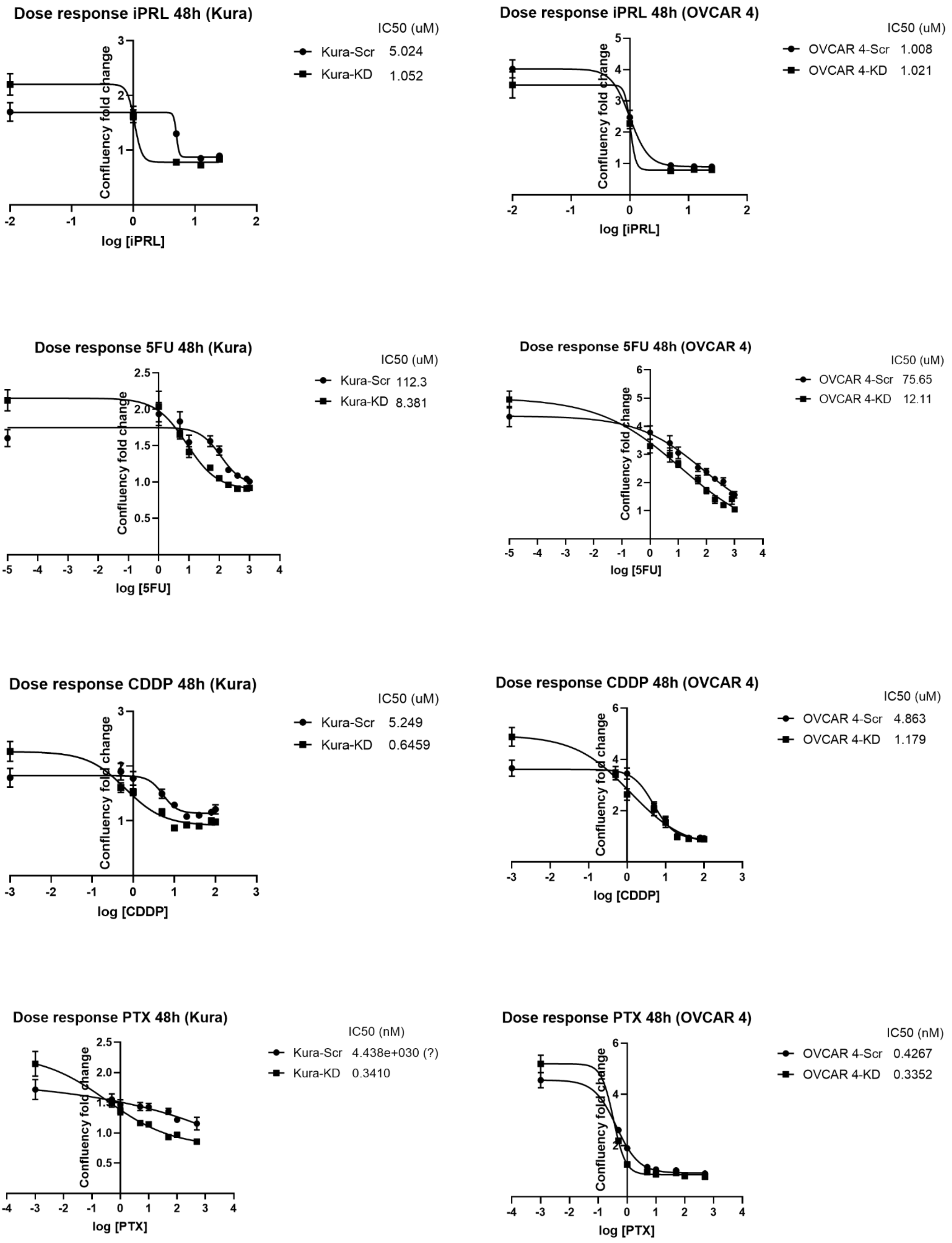
Dose response iPRL 48h (Kura)
IC50 (uM)
Kura-Scr 5.024
Kura-KD 1.052
Confluency fold change
log [iPRL]
Dose response iPRL 48h (OVCAR 4)
IC50 (uM)
OVCAR 4-Scr 1.008
OVCAR 4-KD 1.021
Confluency fold change
log [iPRL]
Dose response 5FU 48h (Kura)
IC50 (uM)
Kura-Scr 112.3
Kura-KD 8.381
Confluency fold change
log [5FU]
Dose response 5FU 48h (OVCAR 4)
IC50 (uM)
OVCAR 4-Scr 75.65
OVCAR 4-KD 12.11
Confluency fold change
log [5FU]
Dose response CDDP 48h (Kura)
IC50 (uM)
Kura-Scr 5.249
Kura-KD 0.6459
Confluency fold change
log [CDDP]
Dose response CDDP 48h (OVCAR 4)
IC50 (uM)
OVCAR 4-Scr 4.863
OVCAR 4-KD 1.179
Confluency fold change
log [CDDP]
Dose response PTX 48h (Kura)
IC50 (nM)
Kura-Scr 4.438e+030 (?)
Kura-KD 0.3410
Confluency fold change
log [PTX]
Dose response PTX 48h (OVCAR 4)
IC50 (nM)
OVCAR 4-Scr 0.4267
OVCAR 4-KD 0.3352
Confluency fold change
log [PTX]

#### Figure S2

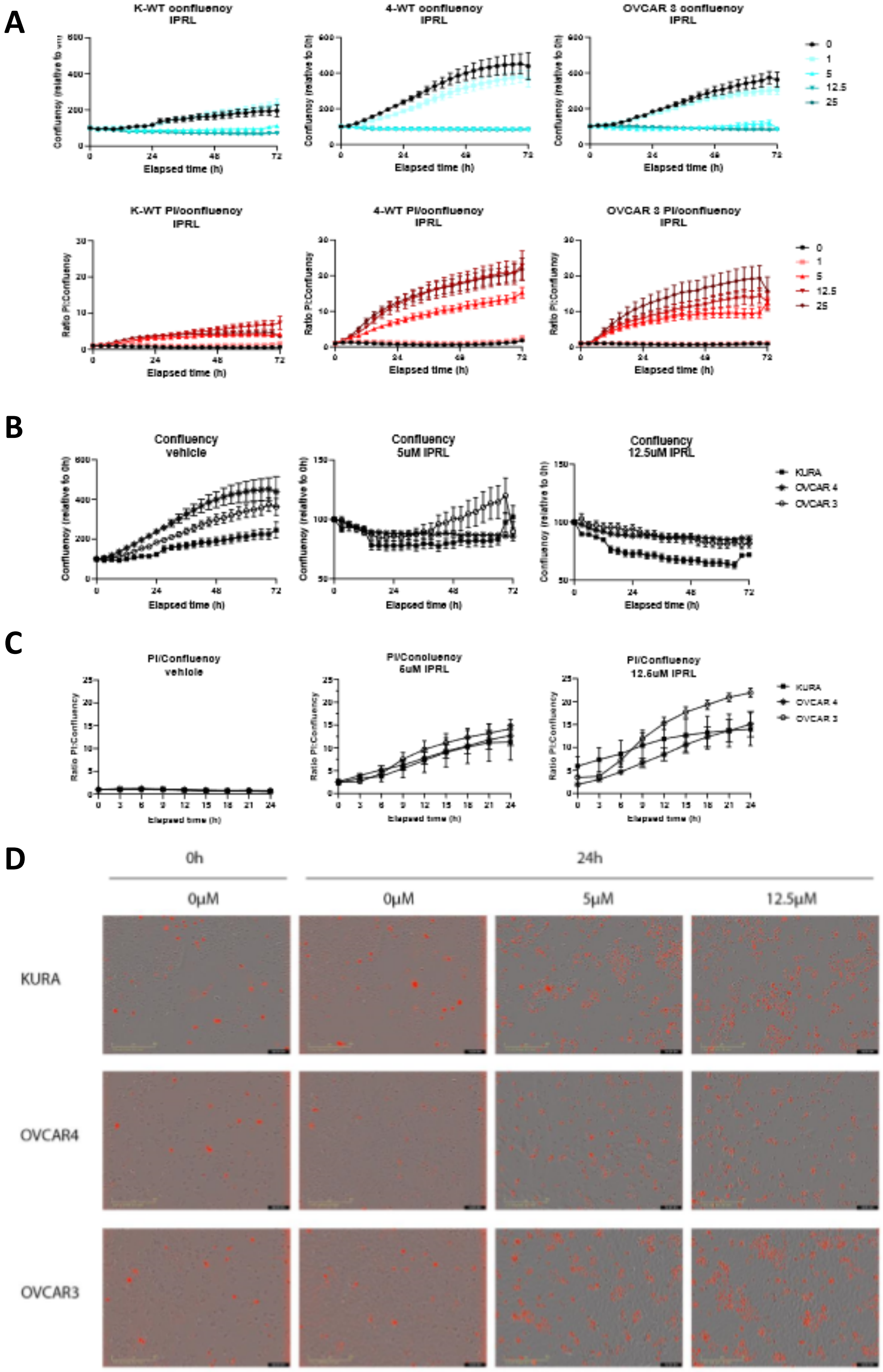
A
K-WT confluency IPRL
4-WT confluency IPRL
OVCAR 3 confluency IPRL
Confluency (relative to 0h)
Elapsed time (h)
0
1
5
12.5
25
K-WT PI/confluency IPRL
4-WT PI/confluency IPRL
OVCAR 3 PI/confluency IPRL
Ratio PI:Confluency
B
Confluency vehicle
Confluency 5uM IPRL
Confluency 12.5uM IPRL
KURA
OVCAR 4
OVCAR 3
C
PI/Confluency vehicle
PI/Confluency 5uM IPRL
PI/Confluency 12.5uM IPRL
D
0h
24h
0µM
5µM
12.5µM
KURA
OVCAR4
OVCAR3

### Figure S3

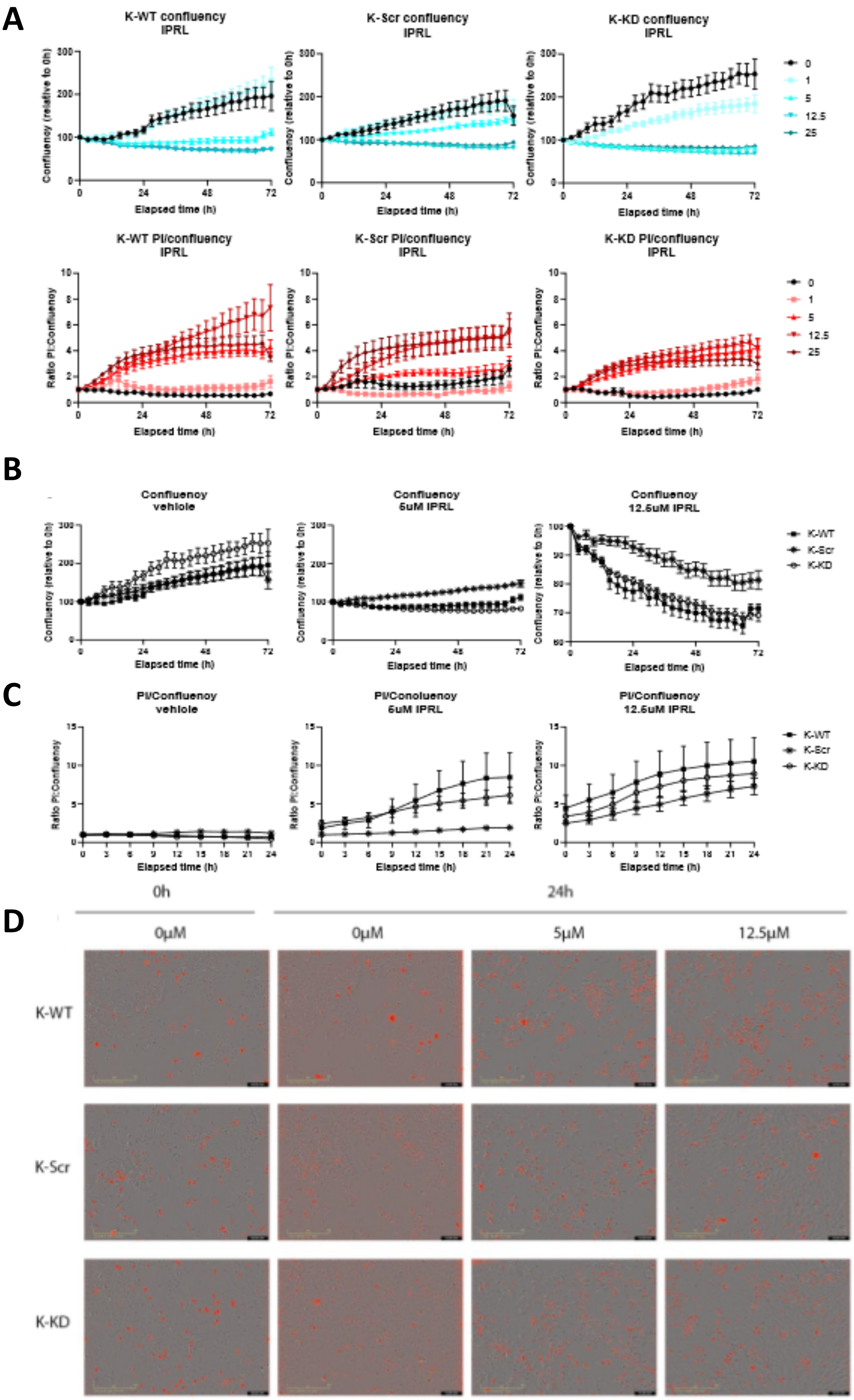

A
K-WT confluency IPRL
K-Scr confluency IPRL
K-KD confluency IPRL
Confluency (relative to 0h)
Elapsed time (h)
0
1
5
12.5
25
K-WT PI/confluency IPRL
K-Scr PI/confluency IPRL
K-KD PI/confluency IPRL
Ratio PI:Confluency
B
Confluency vehicle
Confluency 5uM IPRL
Confluency 12.5uM IPRL
K-WT
K-Scr
K-KD
C
PI/Confluency vehicle
PI/Confluency 5uM IPRL
PI/Confluency 12.5uM IPRL
D
0h
24h
0µM
0µM
5µM
12.5µM
K-WT
K-Scr
K-KD

### Figure S4

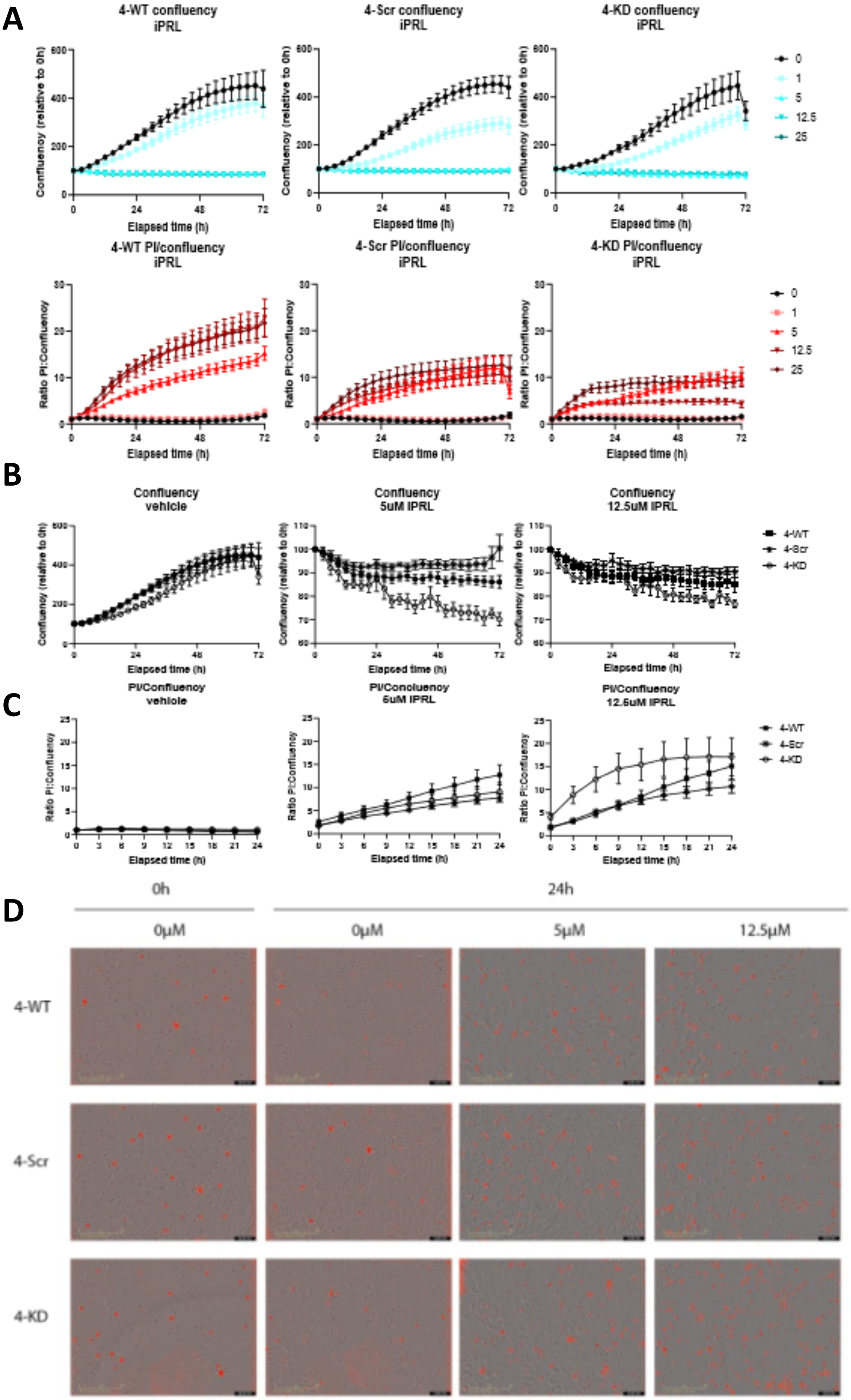

A
4-WT confluency iPRL
4-Scr confluency iPRL
4-KD confluency iPRL
Confluency (relative to 0h)
Elapsed time (h)
0
1
5
12.5
25
4-WT PI/confluency iPRL
4-Scr PI/confluency iPRL
4-KD PI/confluency iPRL
Ratio PI:Confluency
B
Confluency vehicle
Confluency 5uM iPRL
Confluency 12.5uM iPRL
4-WT
4-Scr
4-KD
C
PI/Confluency vehicle
PI/Confluency 5uM iPRL
PI/Confluency 12.5uM iPRL
D
0h
24h
0μM
0μM
5μM
12.5μM
4-WT
4-Scr
4-KD

### Figure S5

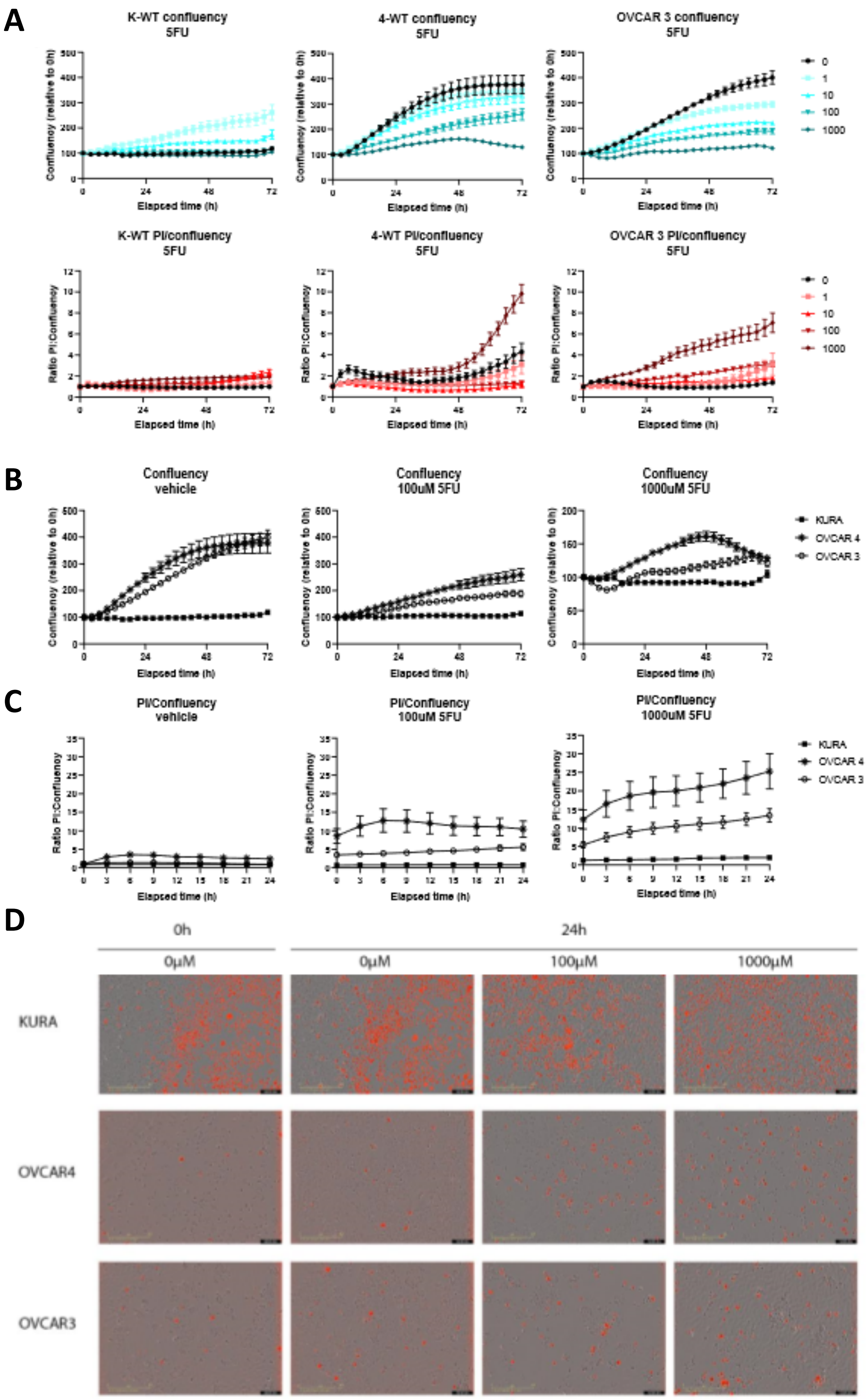
A
K-WT confluency
5FU
4-WT confluency
5FU
OVCAR 3 confluency
5FU
Confluency (relative to 0h)
Elapsed time (h)
0
1
10
100
1000
K-WT PI/confluency
5FU
4-WT PI/confluency
5FU
OVCAR 3 PI/confluency
5FU
Ratio PI:Confluency
B
Confluency
vehicle
Confluency
100uM 5FU
Confluency
1000uM 5FU
KURA
OVCAR 4
OVCAR 3
C
PI/Confluency
vehicle
PI/Confluency
100uM 5FU
PI/Confluency
1000uM 5FU
D
0h
24h
0μM
0μM
100μM
1000μM
KURA
OVCAR4
OVCAR3

### Figure S6

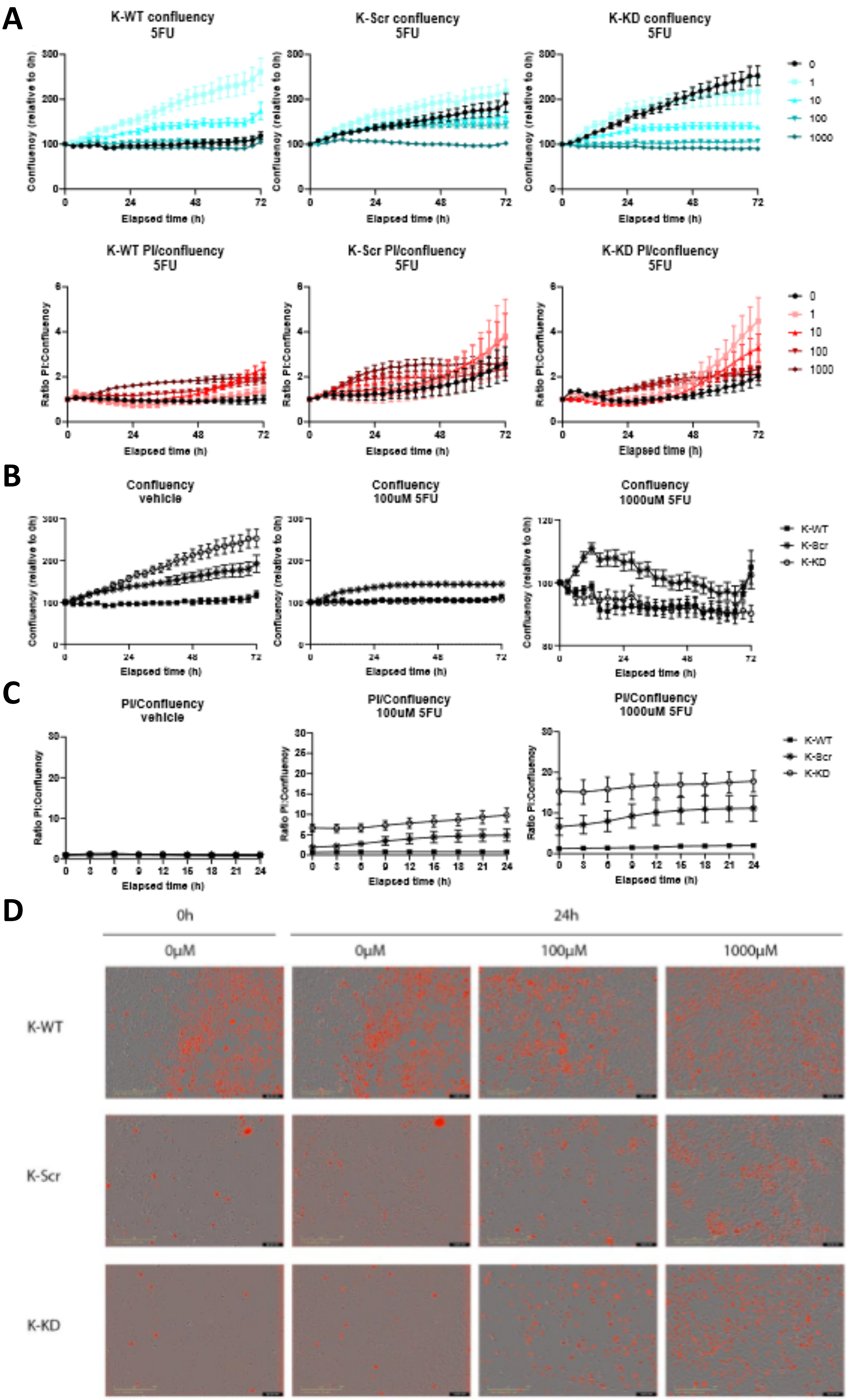
A
K-WT confluency 5FU
K-Scr confluency 5FU
K-KD confluency 5FU
Confluency (relative to 0h)
Elapsed time (h)
0
1
10
100
1000
K-WT PI/confluency 5FU
K-Scr PI/confluency 5FU
K-KD PI/confluency 5FU
Ratio PI:Confluency
B
Confluency vehicle
Confluency 100uM 5FU
Confluency 1000uM 5FU
K-WT
K-Scr
K-KD
C
PI/Confluency vehicle
PI/Confluency 100uM 5FU
PI/Confluency 1000uM 5FU
D
0h
24h
0µM
0µM
100µM
1000µM
K-WT
K-Scr
K-KD

### Figure S7

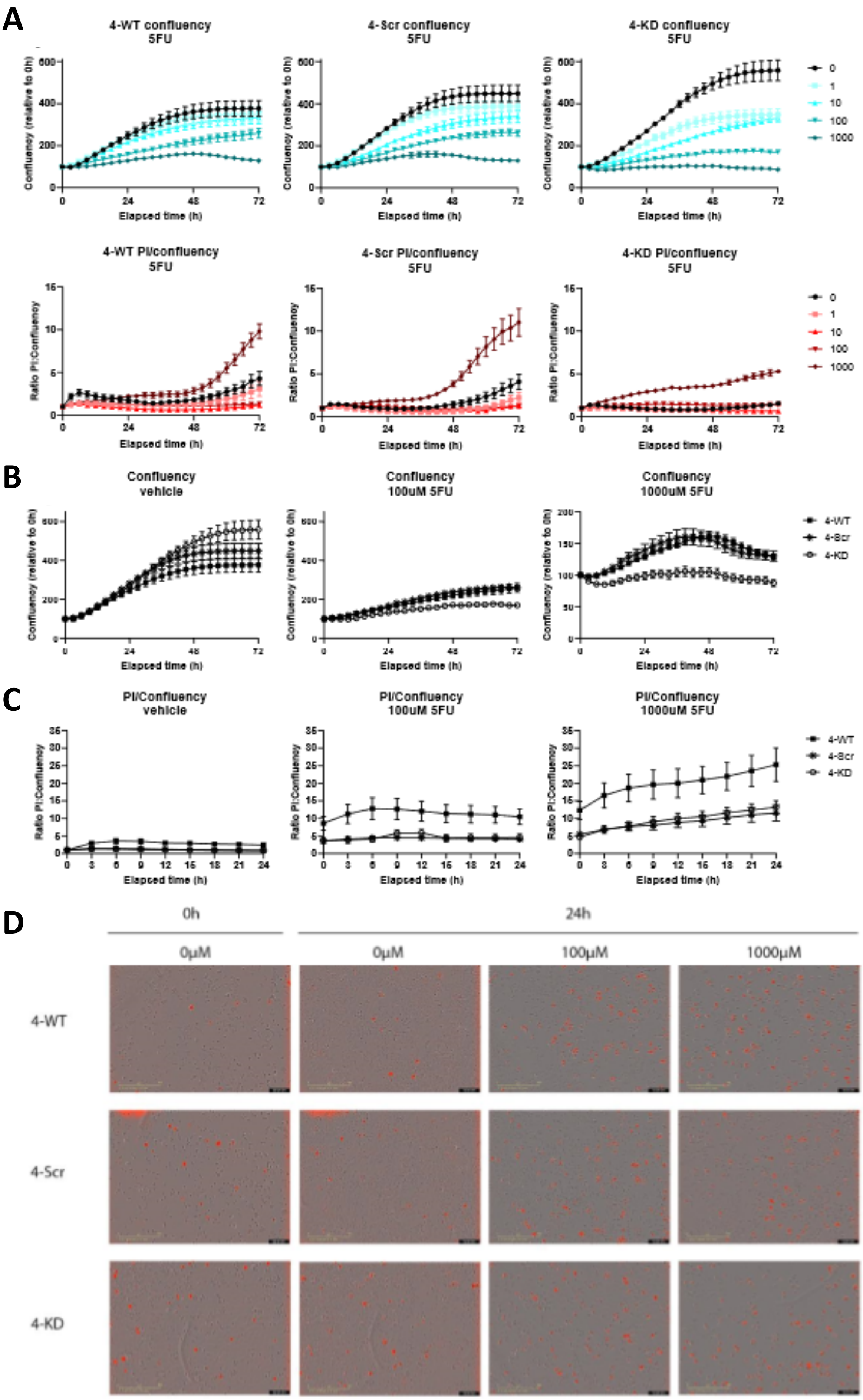
A
4-WT confluency
5FU
4-Scr confluency
5FU
4-KD confluency
5FU
Confluency (relative to 0h)
Elapsed time (h)
0
1
10
100
1000
4-WT PI/confluency
5FU
4-Scr PI/confluency
5FU
4-KD PI/confluency
5FU
Ratio PI:Confluency
B
Confluency
vehicle
Confluency
100uM 5FU
Confluency
1000uM 5FU
4-WT
4-Scr
4-KD
C
PI/Confluency
vehicle
PI/Confluency
100uM 5FU
PI/Confluency
1000uM 5FU
D
0h
24h
0μM
0μM
100μM
1000μM
4-WT
4-Scr
4-KD

### Figure S8

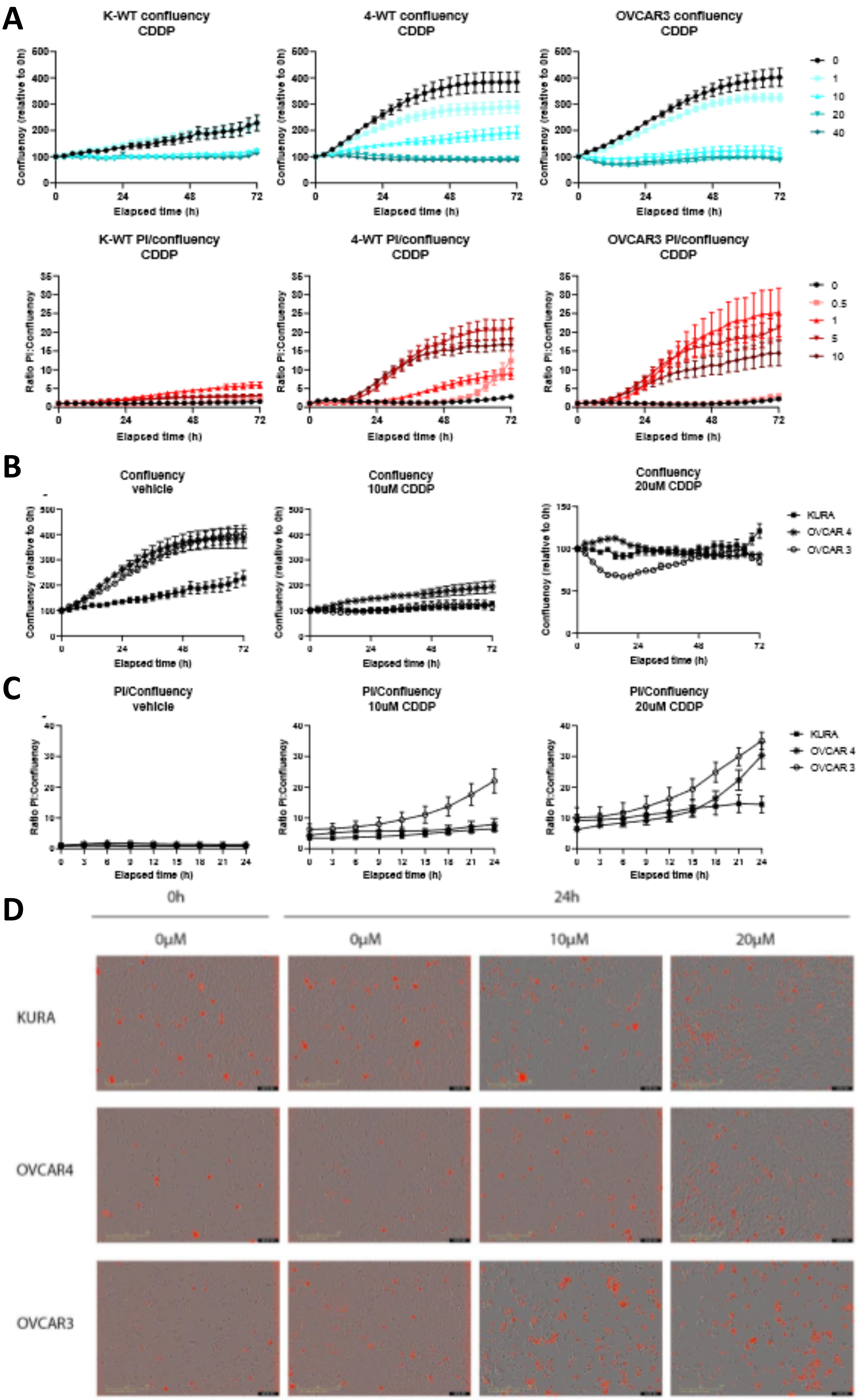
A
K-WT confluency
CDDP
4-WT confluency
CDDP
OVCAR3 confluency
CDDP
Confluency (relative to 0h)
Elapsed time (h)
K-WT PI/confluency
CDDP
4-WT PI/confluency
CDDP
OVCAR3 PI/confluency
CDDP
Ratio PI:Confluency
B
Confluency
vehicle
Confluency
10uM CDDP
Confluency
20uM CDDP
KURA
OVCAR 4
OVCAR 3
C
PI/Confluency
vehicle
PI/Confluency
10uM CDDP
PI/Confluency
20uM CDDP
D
0h
24h
0μM
0μM
10μM
20μM
KURA
OVCAR4
OVCAR3

### Figure S9

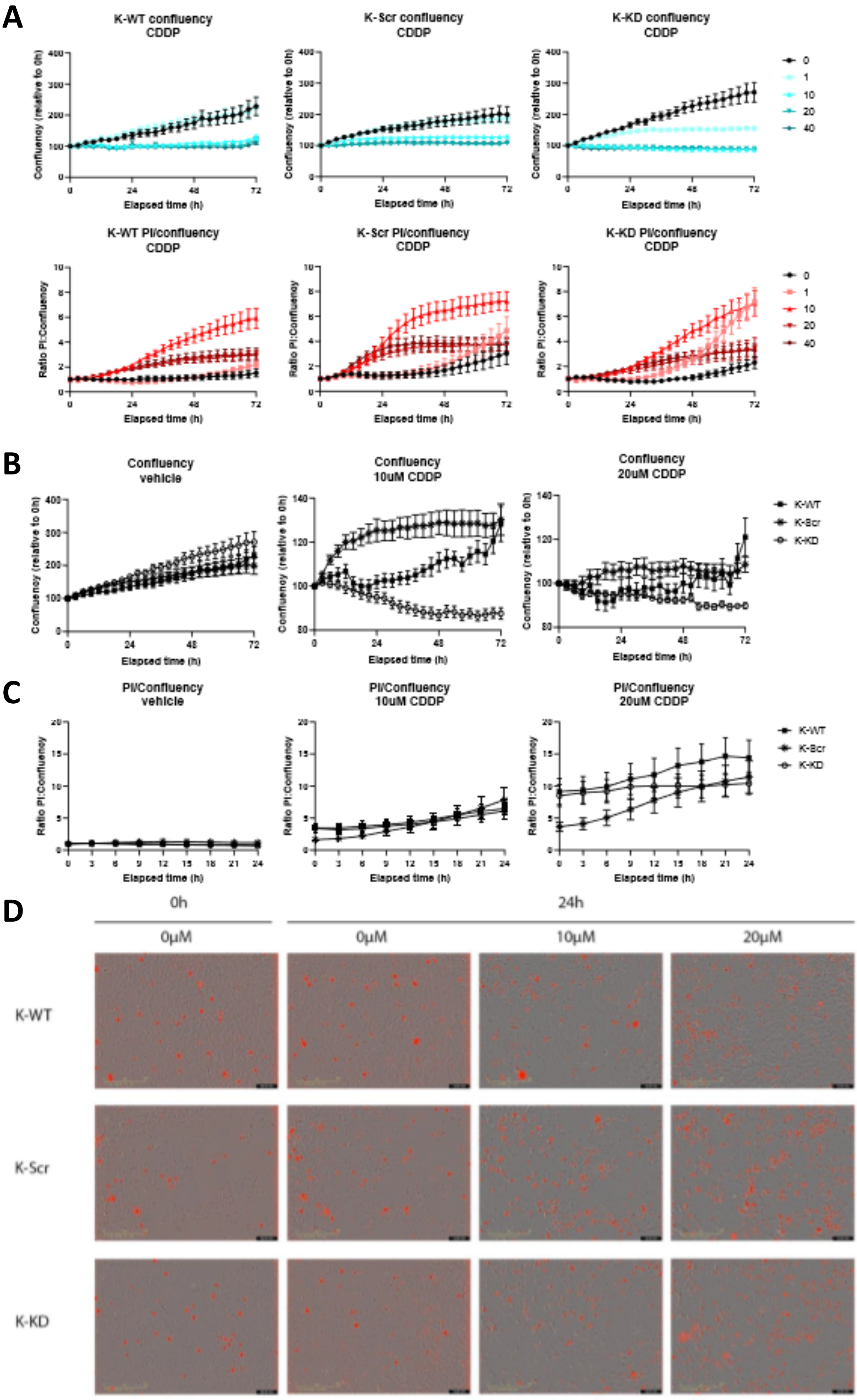

A
K-WT confluency CDDP
K-Scr confluency CDDP
K-KD confluency CDDP
Confluency (relative to 0h)
Elapsed time (h)
0
1
10
20
40
K-WT PI/confluency CDDP
K-Scr PI/confluency CDDP
K-KD PI/confluency CDDP
Ratio PI:Confluency
B
Confluency vehicle
Confluency 10uM CDDP
Confluency 20uM CDDP
K-WT
K-Scr
K-KD
C
PI/Confluency vehicle
PI/Confluency 10uM CDDP
PI/Confluency 20uM CDDP
D
0h
24h
0μM
0μM
10μM
20μM
K-WT
K-Scr
K-KD

### Figure S10

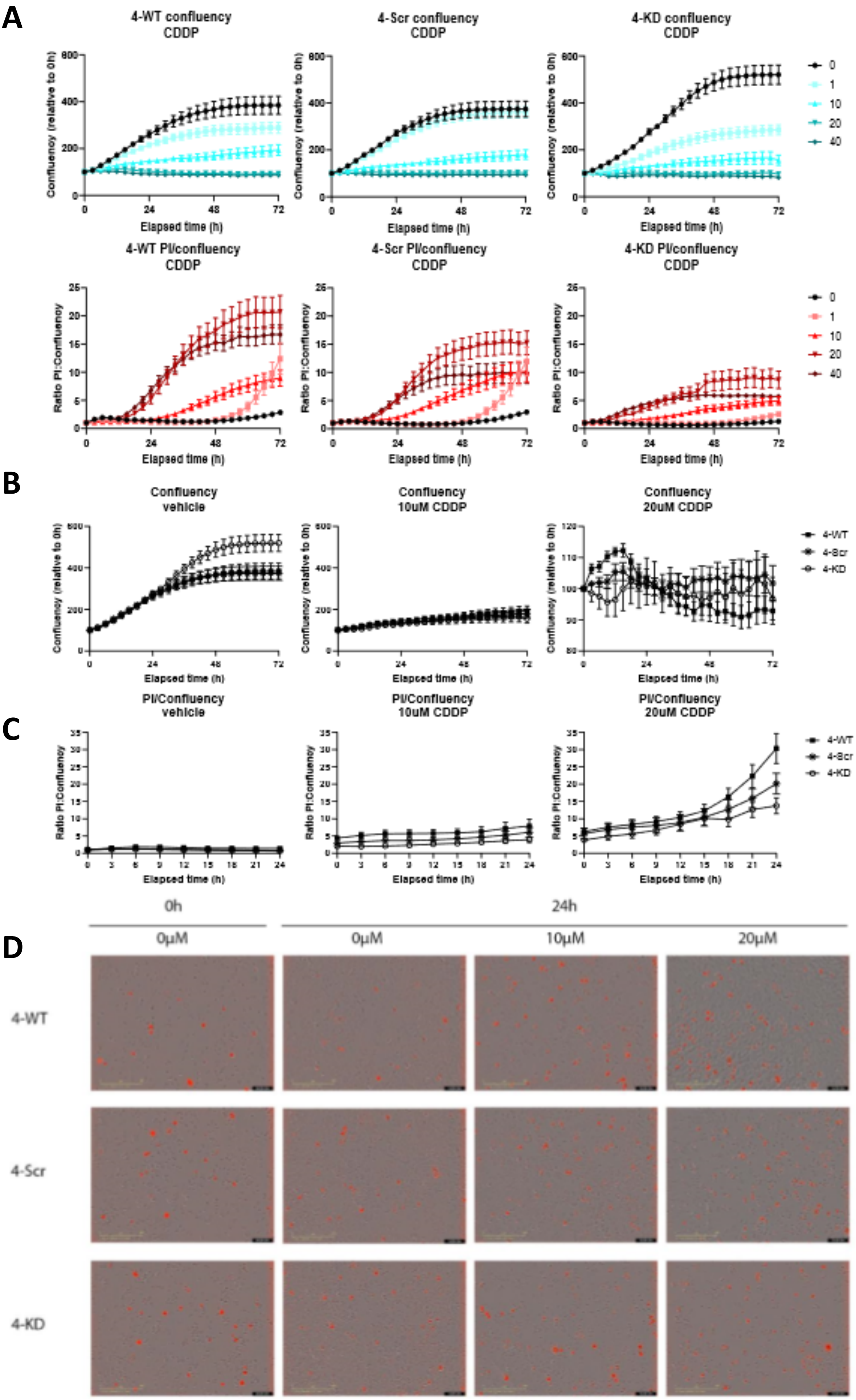
A
4-WT confluency CDDP
4-Scr confluency CDDP
4-KD confluency CDDP
Confluency (relative to 0h)
Elapsed time (h)
0
1
10
20
40
4-WT PI/confluency CDDP
4-Scr PI/confluency CDDP
4-KD PI/confluency CDDP
Ratio PI:Confluency
B
Confluency vehicle
Confluency 10uM CDDP
Confluency 20uM CDDP
4-WT
4-Scr
4-KD
C
PI/Confluency vehicle
PI/Confluency 10uM CDDP
PI/Confluency 20uM CDDP
D
0h
24h
0μM
0μM
10μM
20μM
4-WT
4-Scr
4-KD

### Figure S11

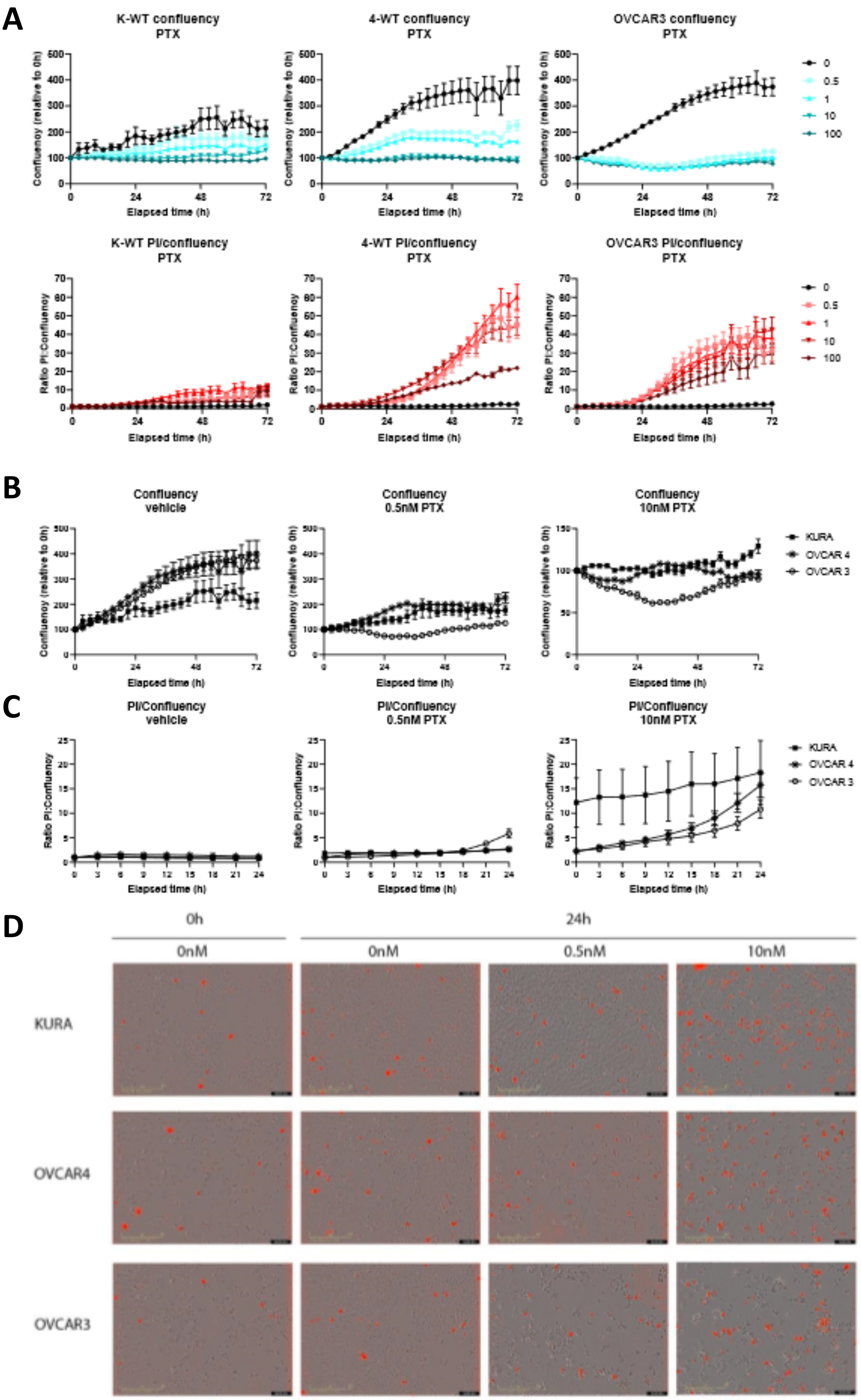
A
K-WT confluency
PTX
4-WT confluency
PTX
OVCAR3 confluency
PTX
Confluency (relative to 0h)
Elapsed time (h)
0
0.5
1
10
100
K-WT PI/confluency
PTX
4-WT PI/confluency
PTX
OVCAR3 PI/confluency
PTX
Ratio PI:Confluency
B
Confluency
vehicle
Confluency
0.5nM PTX
Confluency
10nM PTX
KURA
OVCAR 4
OVCAR 3
C
PI/Confluency
vehicle
PI/Confluency
0.5nM PTX
PI/Confluency
10nM PTX
D
0h
24h
0nM
0nM
0.5nM
10nM
KURA
OVCAR4
OVCAR3

### Figure S12

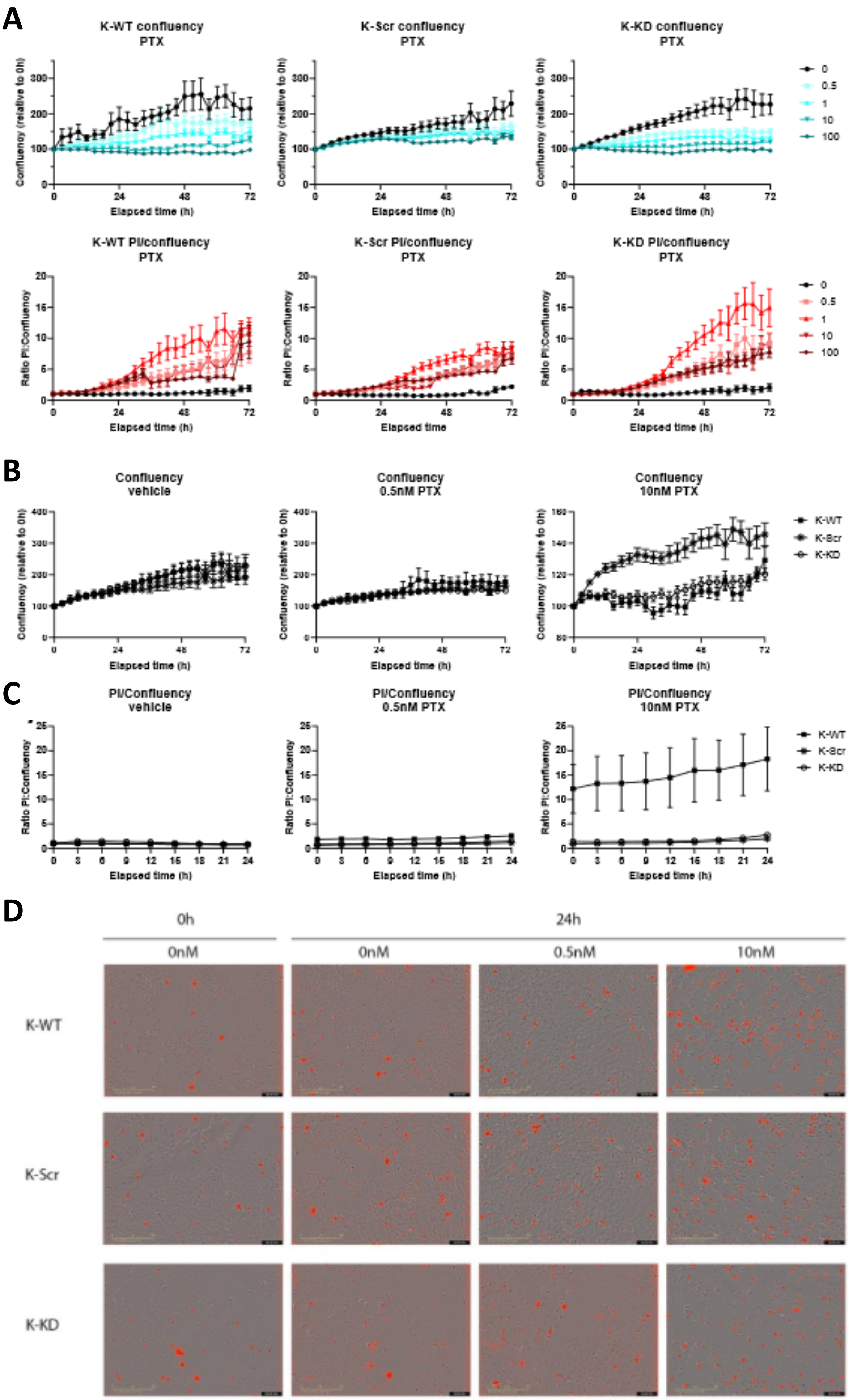
A
K-WT confluency PTX
K-Scr confluency PTX
K-KD confluency PTX
Confluency (relative to 0h)
Elapsed time (h)
0
0.5
1
10
100
K-WT PI/confluency PTX
K-Scr PI/confluency PTX
K-KD PI/confluency PTX
Ratio PI:Confluency
Elapsed time (h)
Elapsed time
B
Confluency vehicle
Confluency 0.5nM PTX
Confluency 10nM PTX
K-WT
K-Scr
K-KD
C
PI/Confluency vehicle
PI/Confluency 0.5nM PTX
PI/Confluency 10nM PTX
D
0h
24h
0nM
0nM
0.5nM
10nM
K-WT
K-Scr
K-KD

#### Figure S13

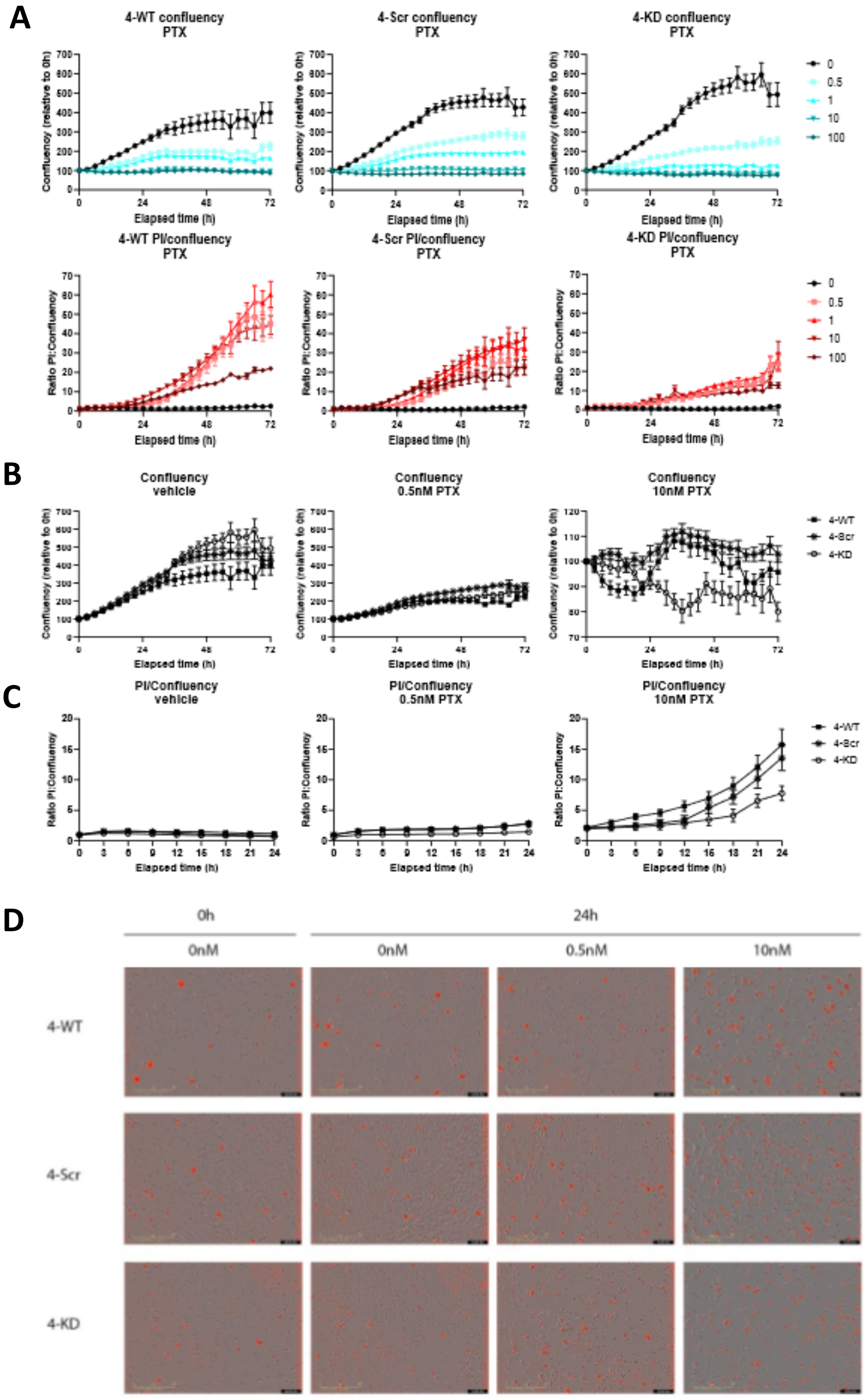
A
4-WT confluency PTX
4-Scr confluency PTX
4-KD confluency PTX
Confluency (relative to 0h)
Elapsed time (h)
0
0.5
1
10
100
4-WT PI/confluency PTX
4-Scr PI/confluency PTX
4-KD PI/confluency PTX
Ratio PI:Confluency
B
Confluency vehicle
Confluency 0.5nM PTX
Confluency 10nM PTX
4-WT
4-Scr
4-KD
C
PI/Confluency vehicle
PI/Confluency 0.5nM PTX
PI/Confluency 10nM PTX
D
0h
24h
0nM
0nM
0.5nM
10nM
4-WT
4-Scr
4-KD

#### Figure S14

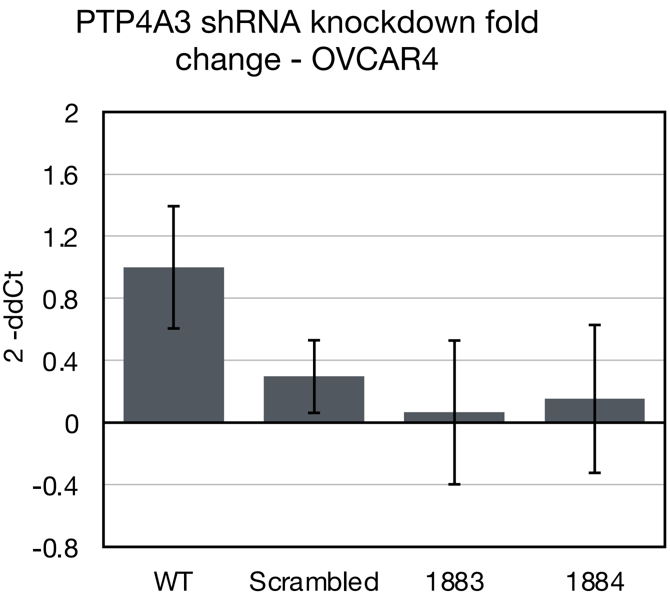

PTP4A3 shRNA knockdown fold change - OVCAR4
2
1.6
1.2
0.8
0.4
0
-0.4
-0.8
2 -ddCt
WT
Scrambled
1883
1884


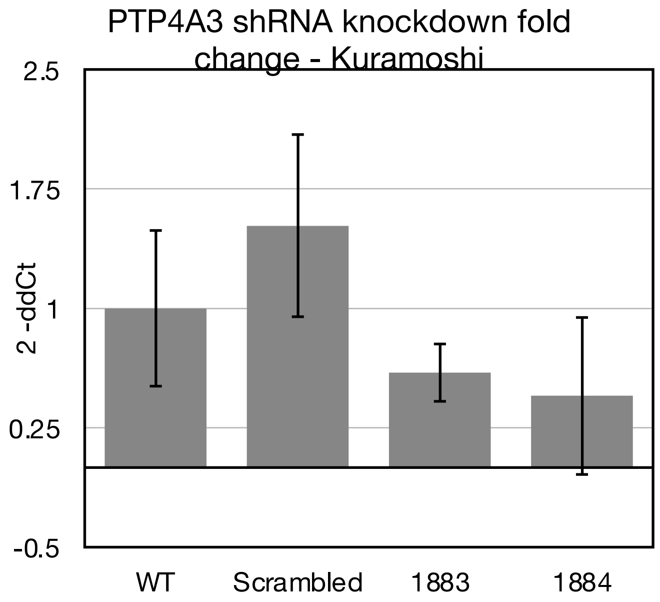

PTP4A3 shRNA knockdown fold change - Kuramoshi
2.5
1.75
1
0.25
-0.5
2 -ddCt
WT
Scrambled
1883
1884
